# Single-cell intracellular epitope and transcript detection revealing signal transduction dynamics

**DOI:** 10.1101/2020.12.02.408120

**Authors:** Francesca Rivello, Erik van Buijtenen, Kinga Matuła, Jessie A.G.L. van Buggenum, Paul Vink, Hans van Eenennaam, Klaas W. Mulder, Wilhelm T. S. Huck

## Abstract

Current high-throughput single-cell multi-omics methods cannot concurrently map changes in (phospho)protein levels and the associated gene expression profiles. We present QuRIE-seq (Quantification of RNA and Intracellular Epitopes by sequencing) and use multi-factor omics analysis (MOFA+) to map signal transduction over multiple timescales. We demonstrate that QuRIE-seq can trace the activation of the B-cell receptor pathway at the minute and hour time-scale and provide insight into the mechanism of action of an inhibitory drug, Ibrutinib.

## Main

The processing of information from the outside environment through signaling pathways is one of the most fundamental processes determining cellular phenotype, function and fate. Single-cell multimodal omics tools are changing our understanding of biology, but we still lack high-throughput single-cell techniques that capture both changes in phosphorylation levels and the much slower ensuing changes in gene expression patterns.^1^ Targeted methods like REAP-seq,^2^ CITE-seq,^3^ or ECCITE-seq,^4^ measure simultaneously transcripts and proteins in single cells using DNA-tagged antibodies, but are limited to the detection of surface epitopes. Our recent work showed a strategy for plate-based single-cell transcriptome and intracellular protein measurements for 6 epitopes.^5^ Here, we present Quantification of RNA and Intracellular Epitopes by sequencing (QuRIE-seq), a high-throughput droplet-based platform to simultaneously quantify 80 intra- and extracellular (phospho)proteins and the transcriptome from thousands of individual cells.

We developed QuRIE-seq to study signal transduction and downstream transcriptional changes in Burkitt lymphoma (BJAB) cells using a validated panel of DNA-barcoded antibodies (Abs) targeting components of the B lymphocyte (B cell) antigen receptor (BCR) signaling pathway, as well as cell cycle, and surface markers (**Supplementary Table 1, Supplementary Fig. 1**). To allow intracellular epitope detection, cells were cross-linked using a mix of reversible fixatives (DSP/SPDP), permeabilized, stained with the DNA-barcoded Ab panel, and co-compartmentalized in nanoliter droplets with single-cell barcoded primer-loaded gel beads (based on the inDrop protocol^6,7^), before further library preparation and sequencing (**Fig. 1a-e**). Reversible fixation with DSP/SPDP and permeabilization of cells showed similar or higher (phospho)protein signals and signal to noise ratios by flow cytometry were equivalent to conventional PFA fixation methods (**Supplementary Fig. 2a,b**). Moreover, the fixation and permeabilization method resulted in a comparable gene-detection rate to unfixed single-cell analysis (**Supplementary Fig. 2c-d**^5^).

**Fig. 1:**
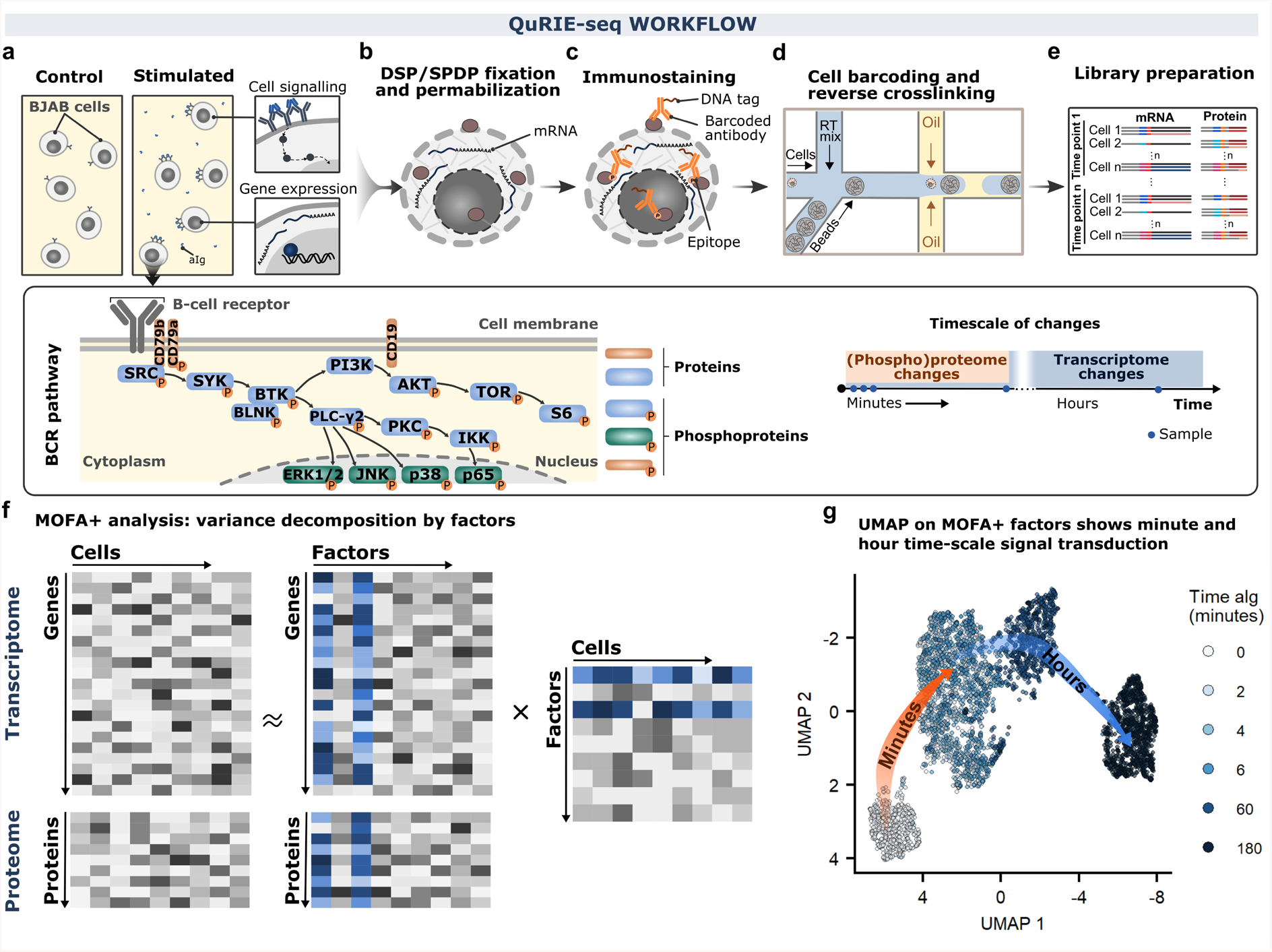
Following multi-modal signal transduction in BJAB cells at the single-cell resolution using Quantification of RNA and Intracellular Epitopes (QuRIE-seq) combined with multi-factor omics analysis (MOFA+) analysis. **a-f**, Schematic overview of the QuRIE-seq workflow for combined single-cell RNA sequencing and (phospho)proteins measurement. **a**, The dynamic response of BJAB cells to stimulation for different durations with polyclonal anti-immunoglobulin antibody (aIg). The scheme of part of the targeted proteins in the B cell antigen receptor signaling (BCR) pathway. The timescale of (phospho)proteome changes fall into minute-regime whereas most significant transcriptome changes are observed after hours. **b**, Cells are reversibly crosslinked with DSP/SPDP (fixatives) and permeabilized. **c**, DNA-tagged antibodies (Abs) targeting membrane and intracellular (phospho)proteins are used for immunostaining. **d**, Single cells are co-encapsulated in a high-throughput manner in nanoliter droplets with barcoding hydrogel beads. **e)** Preparation of two libraries (proteome and transcriptome) is followed by sequencing. **f**, Data analysis overview: MOFA+ takes sequencing data matrices from both modalities (transcriptome and proteome) and decomposes these into matrices containing factors that can be analysed further downstream. **g**, UMAP embedding based on the MOFA+ factors (factor 1 to 7) for 4754 BJAB cells stimulated with aIg for different durations: 0, 2, 4, 6, 60, 180 minutes.

BJAB cells are characterized by functional BCR signaling that can be further stimulated using a polyclonal anti-immunoglobulin antibody (aIg),^8^ as confirmed by aIg induced phosphorylation of components of the BCR signal transduction pathway and aIg concentration-dependent secretion of CCL3 chemokine and IL10, and IL6 cytokines (**Supplementary Fig. 3-4**). We set out to explore molecular changes induced by aIg treatment of BJAB cells at combined (phospho)protein and transcriptome levels across a minute to hour timescale by applying QuRIE-seq on cells stimulated with aIg for 0, 2, 4, 6, 60, and 180 minutes. Data pre-processing included stringent quality control analysis and normalization, resulting in a multimodal dataset of 4754 cells with matched gene and (phospho)protein expression levels. Finally, the data was scaled and corrected for sequencing depth, mitochondrial content (RNA-dataset) and Histone H3 levels (protein dataset) (**Supplementary Fig. 5-6**).

Next, we used unbiased Multi-Omics Factor Analysis (MOFA+)^9^ to combine the quantification of (phospho)proteins and transcripts in a single model that identifies factors explaining variation from both modalities over the full time-series (**Fig. 1f**, **Supplementary Fig. 7**). First, we used the computed MOFA+ factors as input for Uniform Manifold Approximation and Projection (UMAP), capturing the cellular responses to induced activation of the BCR pathway. As shown in **Fig. 1g**, two distinct phases of cellular responses in BJAB cells are observed after short (0-2 minutes) and long (60-180 minutes) stimulation. Exploration of the separate datasets using principal component analysis (PCA) confirmed these observations, showing changes at either minute timescale for the (phospho)protein dataset, and hour timescale for the transcriptome dataset in the first principal component (**Supplementary Fig. 8**), as expected. Together, these initial observations indicate that the computed factors of QuRIE-seq measurements capture the cellular responses to a signal across modalities and timescales.

To explore which molecular changes underlie the BJAB cells response to the aIg stimulus at the minute and hour timescale, we analyzed the computed MOFA+ factors in more detail and found that factors 1 and 3 most strongly correlated with the time of the treatment (**Supplementary Fig. 7c**). Factor 1 captures minute timescale changes (**Fig. 2a**), with predominant effects on the phosphoprotein levels (**Fig. 2b**), as confirmed by flow cytometry (**Supplementary Fig. 9**). Only a minor part of factor 1 is associated with the variance in the mRNA dataset, as transcriptional changes after external stimulation typically take place at the hour, not minute, timescale (**Supplementary Fig. 7a**).

**Fig. 2:**
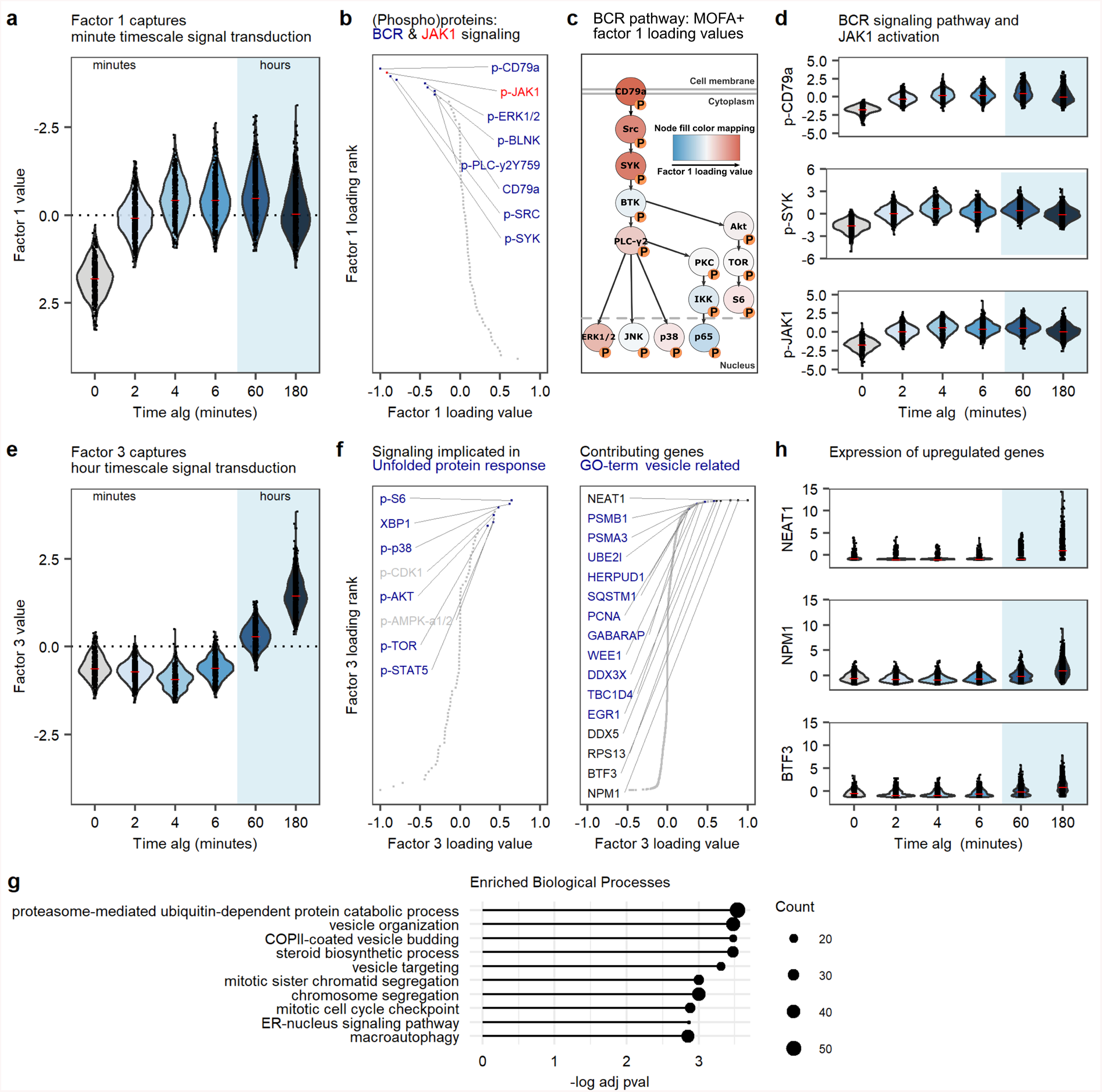
Multi-modal characterization of minute and hour signal transduction. **a**, Violin plot of the factor 1 value as a function of the anti-immunoglobulin antibody (aIg) treatment duration. To reflect the biological intuition on increased phosphorylation upon stimulation, the y-axis of factor 1 is flipped. **b**, Protein loadings contributing to factor 1, ranked according to loading value. Proteins with loading value < −0.25 are annotated in the graphs, colors highlight gene sets related to BCR signaling (blue), or JAK1 (red). **c**, BCR signaling pathway with phospho-proteins colored with MOFA+ factor 1 loading values. **d**, Scaled and normalized QuRIE-seq counts of p-CD79a, p-SYK and p-JAK1 in BJAB cells upon stimulation with aIg. **e**, Violin plot of the factor 3 value as a function of the aIg treatment duration. **f**, Protein (left) and gene (right) loadings contributing to factor 3. Proteins and genes with loading value > 0.25 are indicated in the graphs. Blue color highlights proteins or genes related to indicated biological processes. **g**, Top 10 Gene Ontology (GO) biological process terms enriched in factor 3 genes with positive loading values. **h**, Scaled and normalized QuRIE-seq counts of NEAT1, NPM1 and BTF3 in BJAB cells upon stimulation with aIg.

Examination of the (phospho)proteins and genes with the highest loading values for factor 1 highlighted components of the BCR signaling pathway (e.g. p-CD79a, p-PLC-γ2, p-SRC, and pSYK, *IGHM*, *IGKC*), indicating that factor 1 most likely represents the core elements of the BCR signal transduction pathway, which is activated within two minutes of BCR stimulation with aIg (**Fig. 2c,d, Supplementary Fig. 7 e,f**). Surprisingly, using our unbiased approach p-JAK1 was identified as strongly contributing to factor 1, increasing within the first two minutes after aIg addition, suggesting regulation of JAK1 phosphorylation by BCR activation (**Fig. 2b,d**). Although the Tyrosine kinase JAK1 has been suggested as a substrate for SYK,^10,11^ phosphorylation of JAK1 upon BCR activation has thus far not been described. The unexpected activation of JAK1 by aIg stimulation was further confirmed with flow cytometry (**Supplementary Fig. 10a**). Notably, canonical activation of JAK1 by indirect activation of signaling through cytokines is unlikely to explain these effects, as the observed timescale of two minutes is too short for expression and/or secretion of endocrine cytokines. In addition, we found that the JAK1/2 inhibitor Ruxolitinib resulted in decreased phosphorylation of the BCR pathway components SYK, BTK, PLC-γ2, as well as the JAK1 substrate STAT-6 (**Supplementary Fig. 10b**). This suggests a previously unidentified role for JAK1 in the BCR response to aIg stimulation revealed by QuRIE-seq.

While factor 1 captures changes at minute timescale, factor 3 distinguishes samples at hour timescale (60 and 180 minutes, **Fig. 2e**). The (phospho)proteins with the highest factor 3 loading values are p-S6, XBP1 p-p38, p-AKT, pTOR and pSTAT5 (**Fig. 2f)**. These (phospho)proteins have been shown to contribute to the increase of mis/unfolded proteins in activated B-cells through the Unfolded Protein Response (UPR)^12^, activation of glycolytic energy metabolism and cell cycle entry. RNA features with positive factor 3 loading values (**Fig. 2f,g**) are significantly enriched for gene sets associated with similar cellular function proteasome-mediated protein catabolic process, COPII-mediated vesicle budding, Golgi vesicle-organization and cell-cycle entry. These biological processes relate to increased protein production and energy metabolism, antibody secretion (plasma cell differentiation) and cell-cycle entry (**Fig. 2g, Supplementary Fig. 11**). In addition to proteasome-mediated protein catabolic processes, the protein folding GO-term is enriched (adj. p-value 0.021) in the RNA dataset, consistent with the enrichment of UPR related (phospho)proteins associated with factor 3. The expression of genes with the highest factor 3-loading, including *NEAT1*, *NPM1* and *BTF3*, are upregulated after 60 and 180 minutes of BCR signaling stimulation with aIg (**Fig. 2h**). Comparison of differentially expressed genes determined by bulk RNA-seq and by QuRIE-seq shows similar upregulation profiles at 60 or 180 minutes (**Supplementary Fig. 12**). In summary, the molecular changes at the hour timescale seem to be dominated by proteins and genes involved in processes related to increased protein synthesis, activated glycolysis, B-cell activation and proliferation.

Finally, as QuRIE-seq allows us to map signal transduction at both the (phospho) protein and mRNA level, we explored whether the developed platform can be used to obtain a high-resolution view of the mechanism of action of drugs. Therefore, we stimulated BJAB cells with aIg, as described above, in the presence or absence of 1 μM Ibrutinib (Ibru), a small molecule inhibitor of Bruton’s tyrosine kinase (BTK), blocking a part of the BCR signaling (**Supplementary Fig. 13**). We explored the effect of Ibru treatment at both minute and hour timescale, after 6 and 180 minutes of aIg stimulation. Using MOFA+ we computed a model explaining variation in this dataset (**Supplementary Fig. 14**). UMAP representation shows how aIg stimulation of BJABs in presence of Ibru affects the cellular state at both 6 and 180 minutes time points (**Fig. 3a**). Factor 1 and 3 again correlate with the time of treatment with aIg and capture the short- and long-timescale changes, respectively (**Fig. 3b,c**). Decreased values of factor 1 and 3 for Ibru treated cells (**Fig. 3c**), suggest these processes are, at least partially, BTK dependent. Indeed, phosphorylation of PLC-y2, and BLNK are inhibited by Ibru, as were CD79a, SYK, and JAK1 (**Supplementary Fig. 15a**). The decreased phosphorylation of proteins upstream of BTK suggests the existence of a feedback loop that is affected by Ibru treatment. The expression of the top three genes with high factor 3 loadings shows diminished upregulation after 180 minutes of treatment with aIg (**Supplementary Fig. 15b**). To identify additional effects of BTK inhibition, we note that factor 5 most strongly correlates (Pearson correlation = 0.47) with Ibru treatment and shows increased values after 180 minutes stimulation with aIg for Ibru inhibited cells compared to cells without Ibru inhibition (**Fig. 3b,c**). GO-term analysis shows that features highly contributing to this factor in the positive loadings are two negative regulators of G-coupled protein signaling, *RGS2* and *RGS13* for the RNA-dataset (GO:0045744, adjusted p-value 0.22). Furthermore, in the protein dataset, p-ERK 1/2 contributes, albeit modestly, to factor 5 (**Fig. 3d-f**). Surprisingly, ERK 1/2 phosphorylation is maintained independently of Ibru mediated BTK inhibition (**Fig. 3f**), which supports the notion that Ibru only partially blocks B-cell signal transduction. In line with this, Ibru does indeed inhibit, in a dose-dependent manner, secretion of IL10 and CCL3, but not IL6 (**Supplementary Fig. 16**) supporting the notion of BTK dependent and independent aIg induced activation of BJAB cells. These precursory findings illustrate the potential of QuRIE-seq to study the complexity of inhibitory drug effects on signal transduction.

**Fig. 3:**
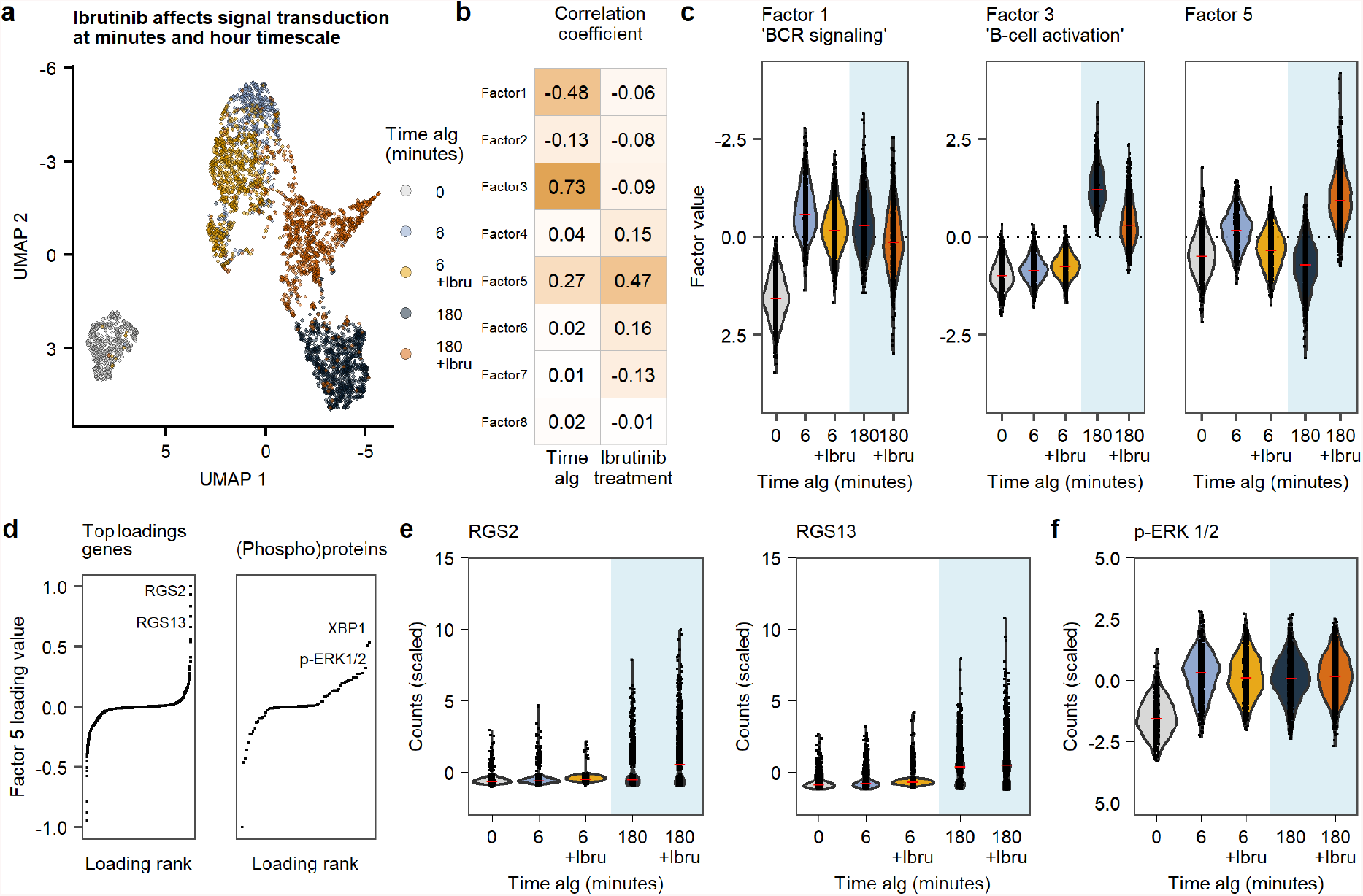
Multi-modal characterization of the dynamic single-cell response to anti-immunoglobulin antibody (aIg) stimulation in presence of Ibrutinib (Ibru) inhibitor of BTK signaling. **a**, UMAP embedding based on the MOFA+ factors for 4658 BJAB cells stimulated with aIg for different durations: 0, 6, 180 minutes with or without Ibru inhibition. **b**, Pearson correlation coefficient by each factor correlating with aIg stimulation duration and with Ibru inhibition. **c**, Violin plots of factor 1, 3, and 5 values as a function of the aIg treatment duration with or without Ibru inhibition. **d**, Gene and protein loadings contributing to factor 5, ranked according to the loading value. **e**, Violin plots of *RGS2* and *RGS13* gene normalized and scaled QuRIE-seq counts. **f**, Violin plot of p-ERK 1/2 normalized and scaled QuRIE-seq counts.

In conclusion, QuRIE-seq combined with MOFA+ analysis of a time series of single-cell intracellular epitope and transcriptome data provides a powerful method to study multimodal intracellular signal transduction at multiple timescales and characterize the mechanism of action of drugs on signal transduction. Presently, we used an antibody panel targeting 80 (phospho)proteins, but we believe there are no biophysical or biochemical limitations to expand this panel to include 500+ target epitopes. Targeting a large number of (phospho)proteins can reveal previously unidentified signaling networks, such as the involvement of JAK1 in BCR signaling or the BTK independent activation of ERK 1/2. Whilst we have exploited droplet-based microfluidics to implement QuRIE-seq, we envision future adaptation to high-throughput plate-based methods such as sci-Plex^13^ and Seq-well.^14^ The combination of customized antibody panels, the ability to detect changes with a high temporal resolution, and the multi-omic read-out of the effect of inhibitory drugs, leads us to anticipate a broad implementation of QuRIE-seq in fundamental signaling studies and drug development research.

## Supporting information

All supplementary information

## Acknowledgments

We thank Sun Tian for assistance with the analysis of RNA bulk sequencing. We acknowledge financial support by Aduro Biotech Europe and Radboud University, Nijmegen.

## Author contributions

W.T.S.H., K.W.M., and H.v.E. supervised the research. F.R., E.v.B., K.M. designed and performed the QuRIE-seq experiment. E.v.B. performed the flow cytometry, ELISA and bulk sequencing experiments. F.R. and J.A.G.L.v.B. analysed the sequencing results. All authors actively contributed to writing and improving the manuscript and approved the final version.

## Experimental methods

### BJAB cells preparation

BJAB cells were obtained from DSMZ, Germany (acc 757). Cells were cultured in RPMI medium (Thermo Fisher Scientific, USA) supplemented with 20% FBS (Biowest, France) and 1% Penicillin-Streptomycin (Thermo Fisher Scientific, USA). Before stimulation, cells were harvested, centrifuged at 200 g for 5 minutes and re-suspended in complete medium at a concentration of 3*10^6^ cells/ml. For each time point, 500 μl cell suspension was dispensed in 3 ml FACS tubes (BD biosciences, USA). For cells treated with inhibitor 1 μM Ibrutinib was added directly to the cell suspension. Subsequently, cells were allowed to rest in complete medium for one hour at 37°C with or without an inhibitor. Cells were then stimulated with 10 μg/ml F(ab’)_2_ Fragment Goat Anti-Human IgA + IgG + IgM (H+L) (Jackson ImmunoResearch, USA) for the indicated time points. After incubation, cells were immediately fixated by adding two times concentrated fixative (5 mM dithiobis(succinimidyl propionate) (DSP) (Thermo Fisher Scientific, USA) and 5 mM succinimidyl 3-(2-pyridyldithio)propionate (SPDP) (Thermo Fisher Scientific, USA) in PBS (Thermo Fisher Scientific, USA), and incubated for one hour at room temperature. Fixated cells were quenched and permeabilized with 100 mM Tris-HCl pH 7.5, 150 mM NaCl and 0.1% Triton X100 (Thermo Fisher Scientific, USA) for 10 minutes at room temperature. After permeabilization, cells were washed once and blocked for 45 minutes in 0.5X protein-free blocking buffer (Thermo Fisher Scientific, USA) with 0.2 mg/ml dextran sulfate (Sigma Aldrich, USA) and 0.5 U/ml RNasin plus (Promega, USA) in PBS (Thermo Fisher Scientific, USA). Cells were stained in a blocking buffer containing DNA-tagged antibodies (Ab list in **Supplementary Table 1**, Abs used for flow cytometry in **Supplementary Table 2**) for one hour at room temperature. Following staining, cells were washed twice with the blocking buffer and re-suspended in PBS containing 5 mg/ml BSA (Thermo Fisher Scientific, USA) and 0.5 U/ml RNsing plus (Promega, USA). Cells were sorted using the BD Biosciences FACS Melody, based on singlets, into protein LoBind tubes (Eppendorf, Germany). Just before encapsulation, each cell suspension was brought to 10^5^ cells/ml and supplemented with 15% (v/v) Optiprep (Sigma Aldrich, USA).

### Microfluidic device fabrication for single-cell barcoding

Microfluidic devices were fabricated by using photo- and soft lithography based on the AutoCAD files with channel designs that were kindly provided by Dr. Linas Mazutisd^5^. Silicon wafers were spin-coated with a uniform layer of SU8-2050 photoresist (MicroChem Co., USA), soft-baked, UV exposed through transparency mask (JD Phototools, UK), baked again post-exposure, developed and hard-baked according to manufacturer’s protocol (MicroChem Co., USA). After production, the height of the structures was measured by Dektak profilometer (the average height ~70 μm). The obtained wafers were used as masters for the PDMS devices. PDMS prepolymer and crosslinking agent were mixed at 10:1 ratio (w/w) and poured on the master bearing the microchannel structure, degassed for 30 minutes and cured at 65 °C for at least 2 hours. Subsequently, the PDMS replica was cut out from the master and the holes were punched at inlet and outlet ports using a biopsy puncher of 1 mm inner diameter (pfm medical, USA). Thereafter, glass slides were carefully washed with a mixture of soap and water, and then both the replicas and the glass slides were rinsed with ethanol. The clean replicas were bonded to the glass slides after oxygen plasma treatment (Femto, Diener electronic, Germany). The channels of the devices were rendered hydrophobic by silanization with 2% 1H,1H,2H,2H-perfluoro-octlytriethoxysilane (Sigma-Aldrich, USA) in FC-40 (Sigma Aldrich, USA). The devices were incubated at 95 °C overnight and afterward stored at room temperature until further usage for droplet production.

### Microfluidic single-cell barcoding

The chip design described by Klein et al.^5^ for single-cell barcoding was used, with four inlets (reverse transcription mix, beads, cell suspension and oil phase with surfactant). The microfluidic single-cell barcoding experimental workflow was followed as described by Zilionis et al^6^ with some modifications. Prior to microfluidic encapsulation, the dispersed and continuous phases used for emulsification were prepared. Barcoded hydrogel beads (1CellBio, USA) were spun down, and mixed with 2X concentration bead mix (0.2% vol/vol Triton X-100 and 2x First Strand buffer) and centrifuged at 5000 g for 2 minutes. The supernatant was removed and the concentrated beads were loaded into the tubing with a previously made air spacer to avoid spreading of beads in HFE-7500 fluorinated fluid (3M) loaded syringe. The cell suspension mixed with 15% (v/v) Optiprep (Sigma-Aldrich, USA) was loaded using a pipette-tip based method for seeding cells to droplet microfluidic platforms, described elsewhere^15^. The reverse transcription (RT) mix comprising 1.3X RT premix MgCl2 (1M), DTT (1M), RNAse OUT (40U/μL), Superscript III (200U/μL) and RNAse free water (Ambion, Thermo Fisher Scientific, USA) was loaded into the tubing. The tubing with the RT mix was wrapped around the ice pack to prevent the increase of temperature and deactivation of enzymes. The oil phase consisted of 3x (v/v) diluted 10% (w/w) RAN (RAN Biotechnologies, USA) in HFE-7500 oil. Solutions were introduced into the microfluidic chip using neMESYS (Cetoni GmbH, Germany) syringe pumps. First, the beads were injected at ~200 μL/h. As soon as the beads appeared at the inlet port, the flow was decreased down to ~20 μL/h, and cell suspension, RT mix, oil spacer were loaded at flow rates ~150 μL/h. When all reagent started to flow through the channels, the final flow rates of the beads, cell suspension, RT mix, oil phase were established ~15-20 μL/h (the flow was tuned during the experiment), 100 μL/h, 100 μL/h, and 90 μL/h, respectively, to produce 2 nl drops. Closely packed barcoding beads were co-encapsulated with single cells in nanolitre droplets, reaching 60-90% cell barcoding efficiency.

After microfluidic emulsification, the cells were de-crosslinked by the action of DTT present in the RT mix which reduced the disulphide bonds present and the barcoding primers were released from the hydrogel beads by 8-minute UV exposure at 10 mW/cm2 from a photolithography machine (ABN Inc, USA). Subsequently, the poly(dT) tail of the barcoding primers annealed both to the poly(A) tail of the mRNA and the poly(dA) tail of the Ab tags and the single cells-derived material was barcoded by reverse transcription: incubation of 2 hours at 50 ℃, followed by 15 minutes at 70 ℃. The emulsion was then broken by adding 3 μL of 20% (vol/vol) PFO on the top of the emulsion, and then 40 μL of HFE-7500 fluorinated fluid (3M). The samples were stored until further use at −80°C.

### Size separation of mRNA and protein libraries

After microfluidic single-cell barcoding, samples were thawed, centrifuged at 4 °C for 5 minutes at 19000 g to pellet cell debris, then the aqueous post-RT material (top layer) was separated from the bottom oil layer and transferred to a nucleic acid purification column (Corning Costar Spin-X column, 0.45 μm, Sigma-Aldrich, USA). Tubes were centrifuged for 1 minute at 16000 g at 4 °C. Subsequently, 100 μL of digestion mix was added per each 70 μL of the sample (final concentrations were 0.5x FastDigest Buffer (Thermo Fisher Scientific, USA) and 0.6 U/μL ExoI (Thermo Fisher Scientific, USA)) and incubated for 30 minutes at 37 °C. After digestion of unused primers and primer dimers, the mix was size separated using 0.6x volume of AMPure XP magnetic beads (Beckman Coulter, USA). The short fragments (the proteomic library that was retained in the supernatant) were separated from the long fragments (the transcriptomic library kept by beads) and each library was then processed independently.

### mRNA library preparation

After the size separation of the mRNA library from the protein library, the mRNA library was processed for second-strand synthesis (SSS) and linear amplification by In Vitro Transcription (IVT). Briefly, 2 μL of 10x SSS buffer and 1 μL of SSS Enzyme (NEBNext mRNA Second Strand Synthesis Module, New England Biolabs, USA) were added to the 17 μL of the purified sample, mixed, and incubated at 16 °C for 1.5 hour, and 20 min at 65 °C. Subsequently, 60 μL of IVT mix was added to each of the 20 μL SSS product, and incubated at 37 °C for 15 hours. The IVT reaction was composed by 1x reaction buffer, 10mM each of ATP, CTP, GTP and UTP, and 10x diluted T7 RNA Polymerase mix (HiScribe™ T7 High Yield RNA Synthesis Kit, New England Biolabs, USA). After linear amplification, the amplified RNA material was purified with 1.3x AMPure XP magnetic beads (Beckman Coulter, USA), and fragmented by adding 1 μL of 10X RNA fragmentation Reagents (Thermo Fisher Scientific, USA) to 10ul of purified IVT, mixing and immediately incubating at 70 °C for exactly 3 minutes. Thereafter, the mix was transferred on ice and 34 μl of STOP mix (8.9 μl nuclease-free water, 24 μl AMPure XP magnetic beads, and 1.1 μl STOP solution) were added to it. Purification using AMPure XP magnetic bead was followed by sample elution in 8 μl nuclease-free water. The following step of the library preparation consisted of the reverse transcription (RT) of the amplified RNA material using random hexamers. This was performed by first addition of 1 μL of 10 nM of each dNTP (Thermo Fisher Scientific, USA) and 2 μL of 100 μM of PE2-N6 primer (Biolegio, The Netherlands), and incubation of the mix at 70 °C for 3 minutes, followed by cooling down the mixture on ice. Next, 3.5 μL nuclease-free water was added to 4 μL of 5x PrimeScript buffer (Takara, USA), 1 μL RNAse OUT (Thermo Fisher Scientific, USA) and 0.5 μL PrimeScript RT (Takara, USA), and the reaction mix was incubated at 30 °C for 10 minutes, followed by 42 °C for 1 hour and by 15 minutes at 70 °C. The RT product was then purified with 1.2x AMPure XP magnetic beads and eluted in 10 μL of nuclease-free water and amplified by PCR. The number of required cycles was determined by qPCR using 0.5 μL of purified material with the addition of 6.5 μL nuclease-free water, 10 μL of 2x Kapa HiFi HotStart PCR mix (Roche, Switzerland), 1 μL of 20x EvaGreen Dye (Thermo Fisher Scientific, USA), and 2 μL of PE1/PE2 primer mix (5 μM each) (Biolegio, The Netherlands). Final material was obtained by mixing 0.5 μL of nuclease-free water, 12.5 μL of 2x Kapa HiFi HotStart PCR mix (Roche, Switzerland), and 2 μL of PE1/PE2 primer mix (5 μM each) (Biolegio, The Netherlands) and the 9.5 μL of the sample. The PCR thermal cycling was as follows: 2 minutes at 98 °C, followed by 2 cycles of: 20 seconds at 98 °C, 30 seconds at 55 °C and 40 seconds at 72 °C, followed by x cycles of (x = number previously determined by qPCR): 20 seconds at 98 °C, 30 seconds at 65 °C and 40 seconds at 72 °C, and terminated with 5 minute-incubation at 72 °C. After PCR, the sample was purified by the addition of 0.7x AMPure XP magnetic beads and eluted in 10 μl of water. The quality of the final libraries was verified by ds-DNA concentration measurement using Qubit ds-DNA High Sensitivity assay (Thermo Fisher Scientific, USA) and BioAnalyzer (Agilent 2100, USA), following the manufacturer’s instructions. If the quality of the samples was sufficient, the mRNA libraries were sequenced together with the protein libraries on with NextSeq500 (targeting 50 million reads per sample). The sequences of the primers used for the library preparations and sequencing are shown in **Supplementary Table 3**.

### Protein libraries preparation

After size separation of the protein and mRNA libraries, the protein library was amplified by IVT reaction. 20 μL of the product was mixed with 60 μL of IVT mix and amplified for 15 hours at 50 °C. The IVT mix was composed of 1x reaction buffer, 10 mM each dNTP (ATP, CTP, GTP and UTP), and 10x diluted T7 RNA Polymerase mix (HiScribe™ T7 High Yield RNA Synthesis Kit, New England Biolabs, USA). After linear amplification, the sample was purified by the addition of 1.5x AMPure XP magnetic beads and eluted in 20 μl of water. 10 μl of the sample were stored at −80 °C as post-IVT backup and the other 10 μl were mixed with 1 μL of 10 nM of each dNTP (Thermo Fisher Scientific, USA), 1 μL of 10 μM PE2-NNNN-Next1 primer (Biolegio, The Netherlands), and 1 μL of 10 μM PE2-NNNN-BioHash2 primer (Biolegio, The Netherlands) and reversely transcribed by incubation for 3 minutes at 70 °C followed by cooling down the mixture on ice. Next, 1.5 μL nuclease-free water, 4 μL of 5x PrimeScript buffer (Takara, USA), 1 μL RNAse OUT (Thermo Fisher Scientific, USA) and 0.5 μL of 200 U/μL PrimeScript RT (Takara, USA) was added to the reaction mix that was incubated: at 30 °C for 10 minutes, followed by 42 °C for 1 hour and by 15 minutes at 70 °C. After reverse transcription, the product was purified with 1.5x AMPure XP magnetic beads and eluted in 10 μL of nuclease-free water. Subsequently, the sample was PCR amplified after prior determination of the number of cycles needed by qPCR: 0.5 μL of purified material was mixed with 6.5 μL nuclease-free water, 10 μL of 2x Kapa HiFi HotStart PCR mix (Roche, Switzerland), 1 μL of 20X EvaGreen Dye (Thermo Fisher Scientific, USA), and 2 μL of PE1/PE2 primer mix (5 μM each) (Biolegio, The Netherlands). Then the PCR reaction was started by adding: 0.5 μL of nuclease-free water, 12.5 μL of 2X Kapa HiFi HotStart PCR mix (Roche, Switzerland), and 2.5 μL of PE1/PE2 primer mix (5 μM each) (Biolegio, The Netherlands) to the 9.5 μL of the sample. The PCR thermal cycling was as follows: 2 minutes at 98 °C, followed by 2 cycles of: 20 seconds at 98 °C, 30 seconds at 55 °C and 40 seconds at 72 °C, followed by x cycles of (x = number previously determined by qPCR): 20 seconds at 98 °C, 30 seconds at 65 °C and 40 seconds at 72 °C, and terminated with 5 minute-incubation at 72 °C. PCR amplified product was purified by addition of 1.2X AMPure XP magnetic beads, eluted in 10 μl water and its quality was verified by ds-DNA concentration measurement with Qubit ds-DNA High Sensitivity assay (Thermo Fisher Scientific, USA) and BioAnalyzer (Agilent 2100, USA) characterization. If the quality of the samples was high enough, the protein libraries were sequenced together with the mRNA libraries on with NextSeq500 (targeting 50 million reads per sample). The sequences of the primers used for the library preparations and sequencing are shown in **Supplementary Table 3**.

### Bioinformatic analysis from FASTQ files to count tables

After sequencing, sequence data from the NextSeq500 (Illumina) was demultiplexed. Two separate FASTQ files were generated for the mRNA and protein libraries. The quality of the sequencing data was evaluated using a FastQC tool (version 0.11.7, Babraham Bioinformatics). The Python script presented by Adrian Veres (https://github.com/indrops/indrops) was used with some modifications, (in particular, for the protein libraries analysis), to process the FASTQ files in order to generate an mRNA and a protein count table. Briefly, the FASTQ files were first filtered to verify that the reads had the correct structure, sufficient quality and complexity. In our setup, 70-80% of the reads were successfully passing the filtering process. Secondly, “real” cells were determined by identifying the cell barcodes having the highest number of reads and setting a threshold below which the barcodes with fewer reads are removed (i.e., cell encapsulated with two beads, a cluster of cells encapsulated with one bead, cells with degraded mRNA). Thirdly, reads were sorted according to their barcode of origin. Subsequently, the remaining reads were aligned using Bowtie with different settings for the mRNA and protein files. For the mRNA m = 10, n = 1, l = 15, e = 1000 as described by Adrian Veres and for the proteins m = 1, n = 1, l = 8, e = 75. With <u>m</u> = the maximum number of different alignments allowed per read, <u>n</u> = the number of mismatches allowed in the first <u>l</u> bases of the read and <u>e</u> = the maximum sum allowed of the quality values al all mismatched positions. The mRNA reads were aligned to the Homo Sapiens Genome FASTA file, while the protein reads were aligned to a self-written FASTA file in which were present the 8 bp Ab tag sequences. Finally, the reads were quantified to UMIFM (UMI-filtered mapped) counts and were aggregated in two separate count tables for the mRNA and for the proteins.

### Quality control, filtering and normalization

To obtain a dataset with high-quality cells several quality checks were performed. First, a Seurat^16^ data object was created, keeping cells with at least 100 genes detected, including genes that were detected in at least 100 cells across the full dataset. Cells with matching protein measurements were added as a second modality, filtering out cells that did appear in only one of the two modalities. Then, cells were filtered out with more than 4000 RNA UMIFM counts and/or less than 150 genes detected, and with more than 15% mitochondrial counts (**Supplementary Fig. 6a-c**). In addition, we filtered out cells with more than 3000 protein counts and/or less than 45 proteins detected (**Supplementary Fig. 6e,f**). The final dataset used for the further analysis, contained 6952 cells, divided over 8 samples: 648 cells at t= 0 minutes, 923 cells at t= 2 minutes, 508 cells at t= 4 minutes, 713 cells at t= 6 minutes, 863 cells at t= 60 minutes, 1099 cells at t= 180 minutes, 943 cells at t= 6 minutes + Ibrutinib, and 1255 cells at t= 180 minutes + Ibrutinib. Next, RNA counts were normalized using ‘single-cell transform’ implemented in the Seurat package^17^, and a scaled dataset was computed where variance in percentage of mitochondrial counts and total UMIFM counts was regressed out. Protein counts were normalized using the CLR method implemented in the Seurat package^16^, and a scaled dataset was computed where variance in number of proteins detected, total UMIFM counts and percentage of Histone H3 counts was regressed out.

### Multi-omics Factor Analysis + (MOFA+) Model on time series of aIg stimulation

Two views were used to train a MOFA (as implemented in MOFA+,^9^) including cells from time-points 0, 2, 4, 6, 60 and 180 minutes with aIg: RNA view included normalized and scaled counts of 2159 variable genes across these samples and the protein view included normalized and scaled counts from all 80 measured (phospho)proteins. MOFA+ default settings were used, resulting in 9 factors. A UMAP was computed on factors 1 to 7. Visualization of the B-cell signaling pathway activity (**Fig. 2c**) with Factor 1 loadings as color scale was done using Cytoscape^18^. Enrichment analysis (using EnrichGO from clusterprofiler^19^) was used to interpret positive gene loadings of factor 3 (**Fig. 2f**).

### Multi-omics Factor Analysis + (MOFA+) Model on timepoints including Ibrutinib data

A MOFA model was built using timepoints 0, 6 and 180 minutes aIg stimulated cells together with 6 and 180 minutes aIg stimulated cells in presence of Ibrutinib. The RNA view included normalized and scaled counts of 2263 variable genes across these samples, and the protein view included normalized and scaled counts from all 80 measured (phospho)proteins. MOFA+ default settings were used, resulting in 8 factors. A UMAP was built on these factors. Loadings (gene and protein dataset) were used to interpret the factors.

### Data and code availability

The data discussed in this publication have been deposited in NCBI’s Gene Expression Omnibus (Edgar *et al*., 2002) and are accessible through GEO Series accession number GSE162461 (https://www.ncbi.nlm.nih.gov/geo/query/acc.cgi?acc=GSE162461).

The code to perform the filtering, analysis and generate the figures in this study is available from Github: https://github.com/jessievb/QuRIE-seq_manuscript.

## References

1. Zhu, C., Preissl, S. & Ren, B. Single-cell multimodal omics: the power of many. Nature Methods 17, 11–14 (2020).

2. Peterson, V. M. et al. Multiplexed quantification of proteins and transcripts in single cells. Nature Biotechnology 35, 936–939 (2017).

3. Stoeckius, M. et al. Simultaneous epitope and transcriptome measurement in single cells. Nature Methods 14, 865–868 (2017).

4. Mimitou, E. P. et al. Multiplexed detection of proteins, transcriptomes, clonotypes and CRISPR perturbations in single cells. Nature Methods 16, 409–412 (2019).

5. Gerlach, J. P. et al. Combined quantification of intracellular (phospho-)proteins and transcriptomics from fixed single cells. Scientific Reports 9, 1469 (2019).

6. Klein, A. M. et al. Droplet Barcoding for Single-Cell Transcriptomics Applied to Embryonic Stem Cells. Cell 161, 1187–1201 (2015).

7. Zilionis, R. et al. Single-cell barcoding and sequencing using droplet microfluidics. Nature Protocols 12, 44–73 (2017).

8. Schamel, W. W. A. & Reth, M. Monomeric and Oligomeric Complexes of the B Cell Antigen Receptor. Immunity 13, 5–14 (2000).

9. Argelaguet, R. et al. MOFA+: a statistical framework for comprehensive integration of multi-modal single-cell data. Genome Biology 21, 111 (2020).

10. Satpathy, S. et al. Systems-wide analysis of BCR signalosomes and downstream phosphorylation and ubiquitylation. Molecular Systems Biology 11, 810 (2015).

11. Patterson, H. C. et al. A respiratory chain controlled signal transduction cascade in the mitochondrial intermembrane space mediates hydrogen peroxide signaling. PNAS 112, E5679–E5688 (2015).

12. Gaudette, B. T., Jones, D. D., Bortnick, A., Argon, Y. & Allman, D. mTORC1 coordinates an immediate unfolded protein response-related transcriptome in activated B cells preceding antibody secretion. Nature Communications 11, 723 (2020).

13. Zheng, G. X. Y. et al. Massively parallel digital transcriptional profiling of single cells. Nature Communications 8, 14049 (2017).

14. Gierahn, T. M. et al. Seq-Well: portable, low-cost RNA sequencing of single cells at high throughput. Nature Methods 14, 395–398 (2017).

15. Sinha, N., Subedi, N., Wimmers, F., Soennichsen, M. & Tel, J. A Pipette-Tip Based Method for Seeding Cells to Droplet Microfluidic Platforms. JoVE (Journal of Visualized Experiments) e57848 (2019).

16. Stuart, T. et al. Comprehensive Integration of Single-Cell Data. Cell 177, 1888–1902.e21 (2019).

17. Hafemeister, C. & Satija, R. Normalization and variance stabilization of single-cell RNA-seq data using regularized negative binomial regression. Genome Biology 20, 296 (2019).

18. Shannon, P. et al. Cytoscape: A Software Environment for Integrated Models of Biomolecular Interaction Networks. Genome Res. 13, 2498–2504 (2003).

19. Yu, G., Wang, L.-G., Han, Y. & He, Q.-Y. clusterProfiler: an R Package for Comparing Biological Themes Among Gene Clusters. OMICS: A Journal of Integrative Biology 16, 284–287 (2012).

