## Supplementary material for "Single-cell intracellular epitope and transcript detection revealing signal transduction dynamics": All supplementary information

#### Contents

Supplementary Notes 3

Supplementary Figures 5

Supplementary Tables 22

### Supplementary Notes

#### I. Antibody labeling

All antibodies (Abs) were purchased purified (see **Supplementary Table 1** for a full list of the Abs used). Abs used in the QuRIE-seq experiment were first validated using flow cytometry.

QuRIE-seq requires the use of DNA-tagged Abs for staining cells prior to encapsulation to characterize the single-cell proteome. In the following is briefly detailed the DNA conjugation strategy to the Abs which made us of standard NHS chemistry. First, Abs were buffer exchanged into 0.2M $NaHCO_3$ (pH 8.3) (Sigma Aldrich, USA) using Zeba™ Spin Desalting Columns, 40K MWCO (Thermo Fisher Scientific, USA). Then, dibenzocyclooctyne-S-S-N-hydroxysuccinimidyl ester (Sigma Aldrich, USA) was resuspended in DMSO (Sigma Aldrich, USA), added in a 10x molar excess to the Abs and incubated for 2 hours at room temperature. Abs were then buffer exchanged into PBS (Thermo Fisher Scientific, USA) and excess linker molecules were removed using 30K amicon centrifuge units (Merck, USA). Subsequently, 5' Azide modified oligos (Biolegio, The Netherlands) were added in a 3x molar excess to the chemically modified Abs and incubated for 16 hours at 4°C in dark. After incubation, Abs were buffer exchanged into PBS (Thermo Fisher Scientific, USA) containing 0.05% sodium azide (Sigma Aldrich, USA) and 0.1mM EDTA (Sigma Aldrich, USA), and excess oligos were removed using 100K amicon centrifuge filters (Merck, USA). After labelling, DNA-tagged Abs were stored at 4°C. Labelling efficiency was determined by non-reducing SDS page gel analysis where 0.5 µg antibody was diluted in 1x Laemmli sample buffer (Bio-Rad, USA) and loaded on a mini-PROTEAN TGX stain-free gel (Bio-Rad, USA). Gels were imaged on a ChemiDoc Touch imaging system (Bio-Rad, USA).

#### II. Flow cytometry analysis

DSP and PDSP fixed and permeabilized BJAB cells were centrifuged at 800 g and resuspended in PBS (Thermo Fisher Scientific, USA) supplemented with 0.1% BSA (Thermo Fisher Scientific, USA) containing the indicated Abs, and incubated for one hour at room temperature in dark (**Supplementary Table 2** shows different Abs used for flow cytometry analysis). After incubation, cells were washed three times with PBS containing 0.1% BSA and resuspended in PBS with 0.1% BSA. Finally, cells were measured on a FACSverse (BD Bioscience, USA). Flow cytometry results were analysed using FlowJo software (BD Bioscience, USA).

#### III. ELISA

BJAB cells were cultured in round bottom 96-wells plates at a concentration of $10^5$ cells/well in a volume of 200 µl. Cells were stimulated with 0-20 µg/ml F(ab’) $_2$ Fragment Goat Anti-Human IgA + IgG + IgM (H+L) (Jackson ImmunoResearch, USA) and incubated for 16 hours at 37 °C. The supernatant was collected and analysed for the presence of IL6, IL10 and CCL3 using uncoated ELISA kit (Thermo Fisher Scientific, USA). ELISA was performed according to the manufacturer's instructions.

#### IV. Bulk mRNA Sequencing

BJAB cells were harvested, centrifuged and resuspended in complete medium at $0.5*10^6$ cells/ml. Thereafter, 4 ml of cells were seeded in T25 flasks (Thermo Fisher Scientific, USA), and cells were incubated for 1 hour at 37 °C. After incubation, the cells were stimulated with 10 µg/ml F(ab')$_2$ Fragment Goat Anti-Human IgA + IgG + IgM (H+L) (Jackson ImmunoResearch, USA) for the indicated time points. After stimulation, cells were immediately collected on ice and centrifuged at 200 g for 5 minutes at 4°C. Supernatant was collected and stored at -20°C for further analysis and the cell pellet was resuspended in the lysis buffer RLT (Qiagen, The Netherlands). Cell lysates were stored at -80°C until further use. Library preparation and sequencing was performed by Macrogen (Macrogen, Korea).

#### V. Bulk mRNA Sequencing data analysis

RNAseq fastq files of samples were aligned to the human genome GRCh38 using HISAT2 v2.1.0. Number of reads was assigned to genes by using featureCounts v1.6.1. Reads mapped to genes were normalized using count per million method (CPM) implemented in edgeR package in R Bioconductor. To analyze genes that have the highest fold change upon different durations of stimulation with aIg (60, and 180 min), genes were ranked by the difference between their CPM normalized values and mean of CPM normalized values of this gene in the unstimulated control sample.

#### Supplementary Figures

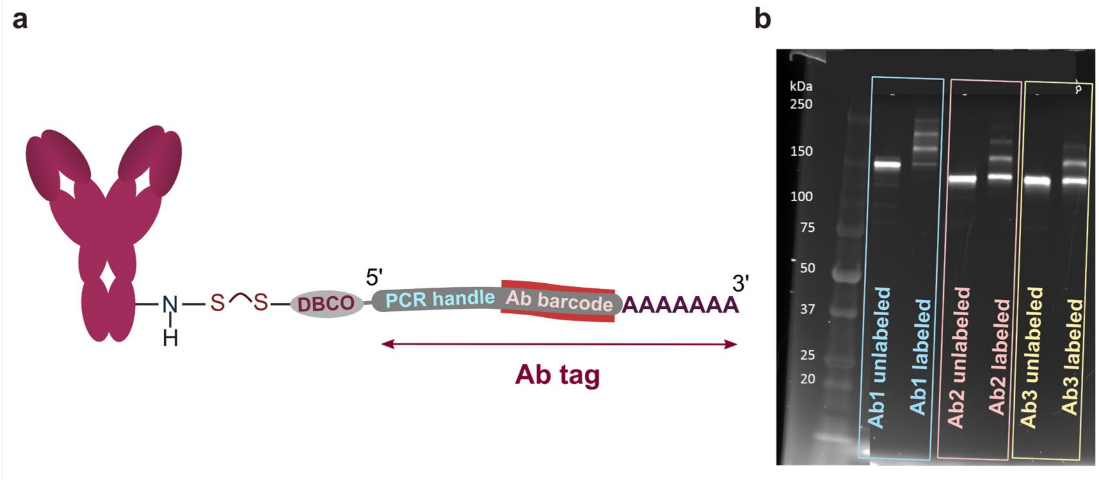


**Supplementary Figure 1: Antibodies (Abs) labelling for single-cell proteome quantification in Quantification of RNA and Intracellular Epitopes (QuRIE-seq). a**, Scheme of the Ab with a DNA tag. *DBCO = Dibenzocyclooctyne.* **b**, SDS page gel for the determination of Ab labelling efficiency where 0.5 µg Ab was diluted in non-reducing 1x Laemmli sample buffer (Bio-Rad) and loaded on a mini-PROTEAN TGX stain-free gel (Bio-Rad). Gels were imaged on a ChemiDoc Tough imaging system (Bio-Rad).

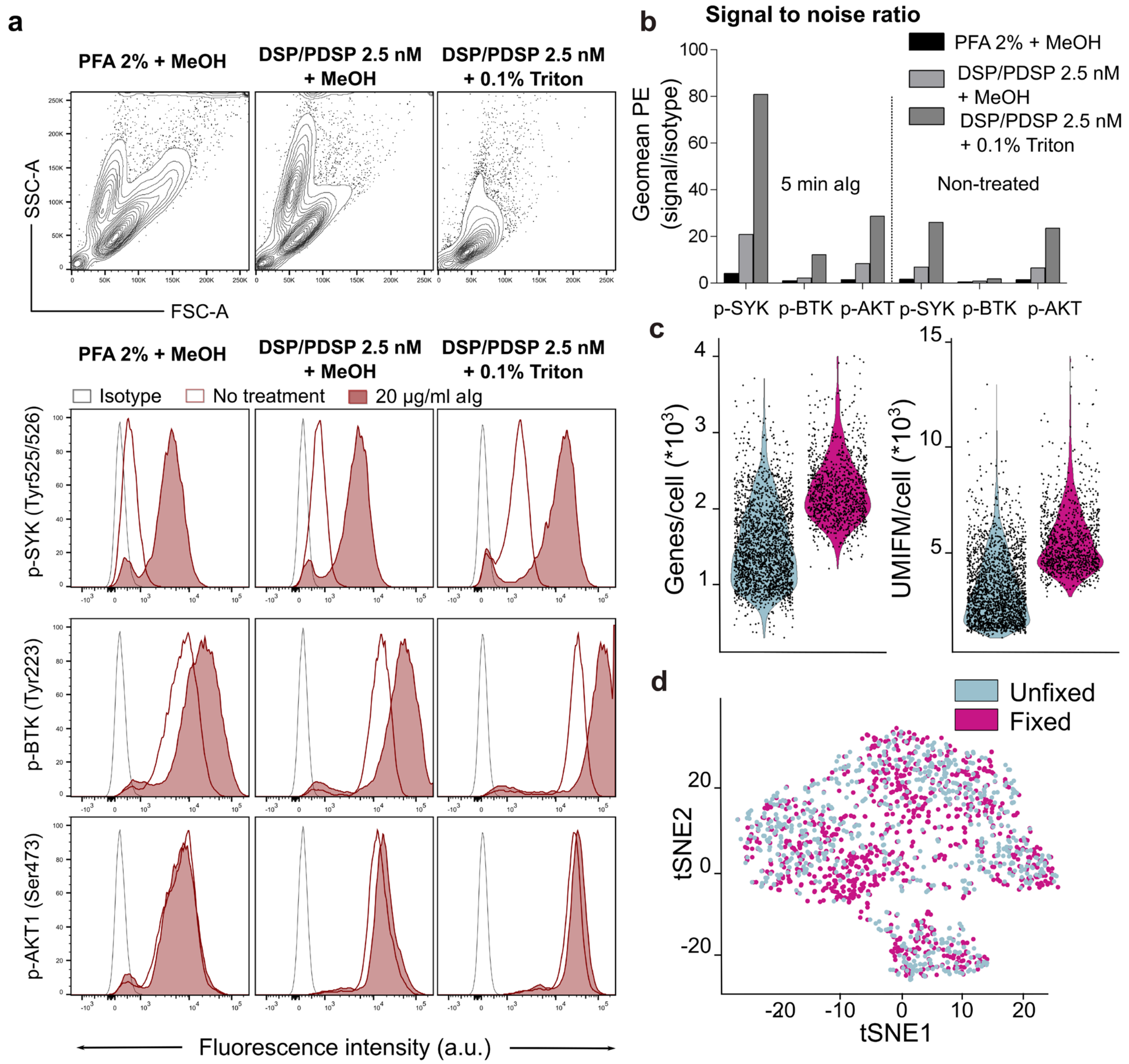


**Supplementary Figure 2: Effect of the fixation procedure with BJAB cells on antibody binding, detection of phosphorylated proteins, and gene detection. a**, Two different fixation methods were compared with flow cytometry: DSP/PDSP and PFA fixation both in combination with methanol (MeOH) permeabilization, and two permeabilization methods: MeOH and Triton-X100 in combination with DSP/PDSP fixation. The phosphorylation of SYK, BTK and AKT after 5 minute-stimulation with αIg. **b**, The signal to noise ratio was compared with flow cytometry for two different fixation methods: DSP/PDSP and PFA in combination with MeOH permeabilization, and two permeabilization methods: MeOH and Triton-X100 in combination with DSP/PDSP fixation. **c**, Comparison of the number of genes and the number of UMIFM (UMI-filtered mapped) detected per cell between unfixed and fixed BJABs using QuRIE-seq technology. **d**, t-SNE visualization of the unfixed and fixed BJABs transcriptome expression using QuRIE-seq technology.

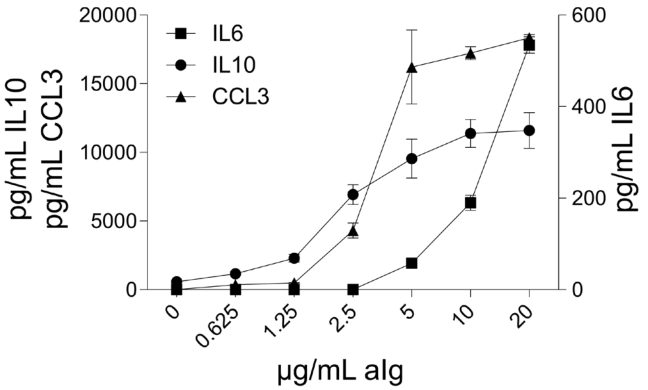


**Supplementary Figure 3: Analysis of BJABs culture supernatant for cytokines upon overnight stimulation with polyclonal anti-immunoglobulin antibody (aIg).** BJAB cells were stimulated for 16 hours with a titration of aIg after which the supernatant was analysed by ELISA for IL6, IL10 and CCL3.

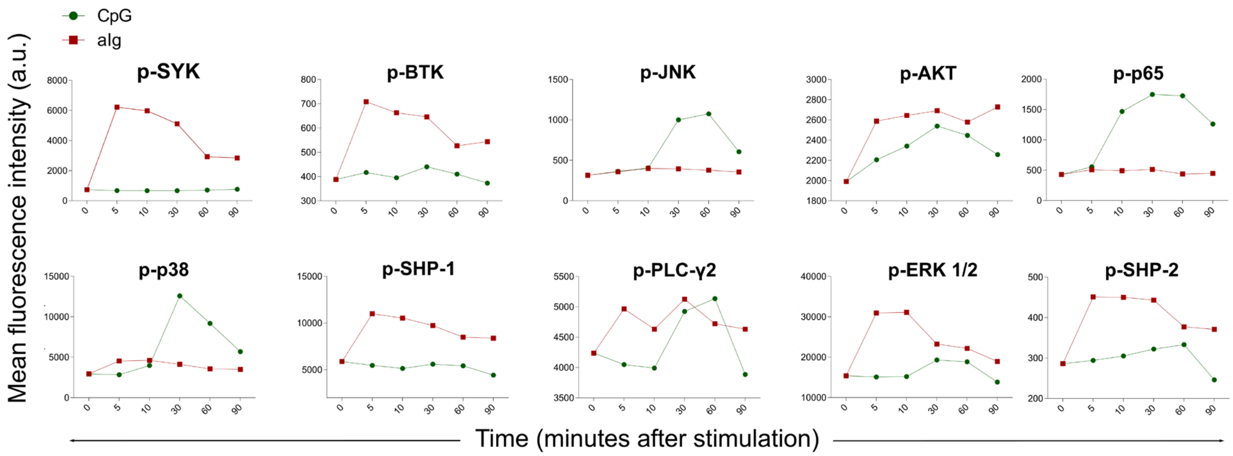


**Supplementary Figure 4: Validation by flow cytometry of stimulation by polyclonal anti-immunoglobulin antibody (aIg) of the BCR pathway.** The phosphorylation of SYK, BTK, JNK, AKT, p65, p38, SHP-1, PLC-γ2, ERK 1/2, and SHP-2 was characterized upon different stimulation conditions: unstimulated cells (0 minutes); stimulation with aIg (red line) or CpG (green line) for: 5, 10, 30, 60 and 90 minutes. *CpG stimulation targeting the toll-like receptor (TLR9) was used as a control since it is expected to have different dynamics compared to aIg.*

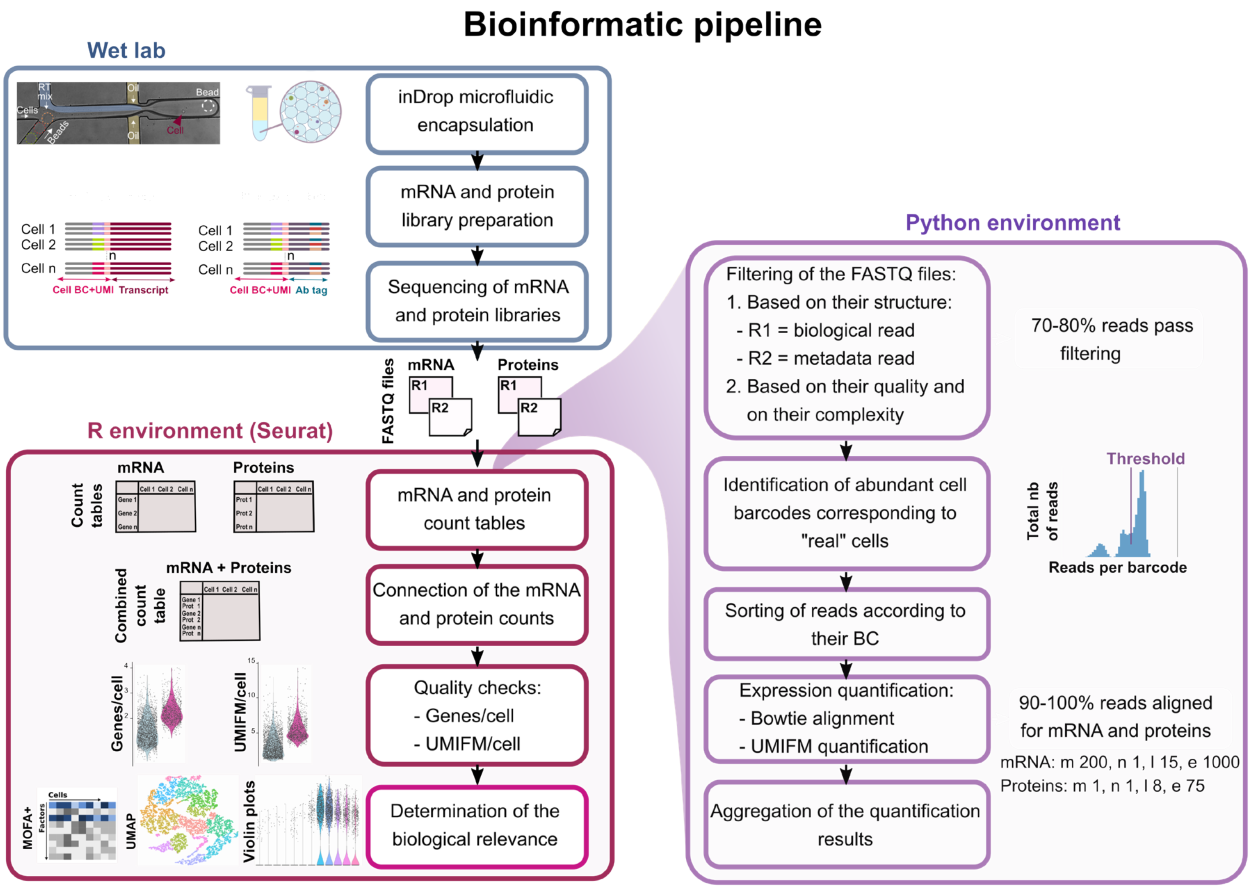


**Supplementary Figure 5: Workflow and extended bioinformatics pipeline for the analysis of results obtained with QuRIE-seq platform. In the wet lab**, cells are encapsulated together with barcoding beads using QuRIE-seq technology. Subsequently, an mRNA and a protein library are prepared in parallel and sequenced. **In the Python environment**, the FASTQ files are processed to generate mRNA and protein count tables. This processing comprehends: filtering of the reads, determination of the "real" cells, mapping of the reads to the reference genome using bowtie alignment, and quantification of UMIFM (UMI-filtered mapped) counts. With m = the maximum number of different alignments allowed per read, n = the number of mismatches allowed in the first l bases of the read and e = the maximum sum allowed of the quality values at all mismatched positions. **In the R environment**, the two mRNA and protein count tables are connected in one combined count table. These count tables are the basis for quality check, normalization and scaling, and MOFA+ analysis.

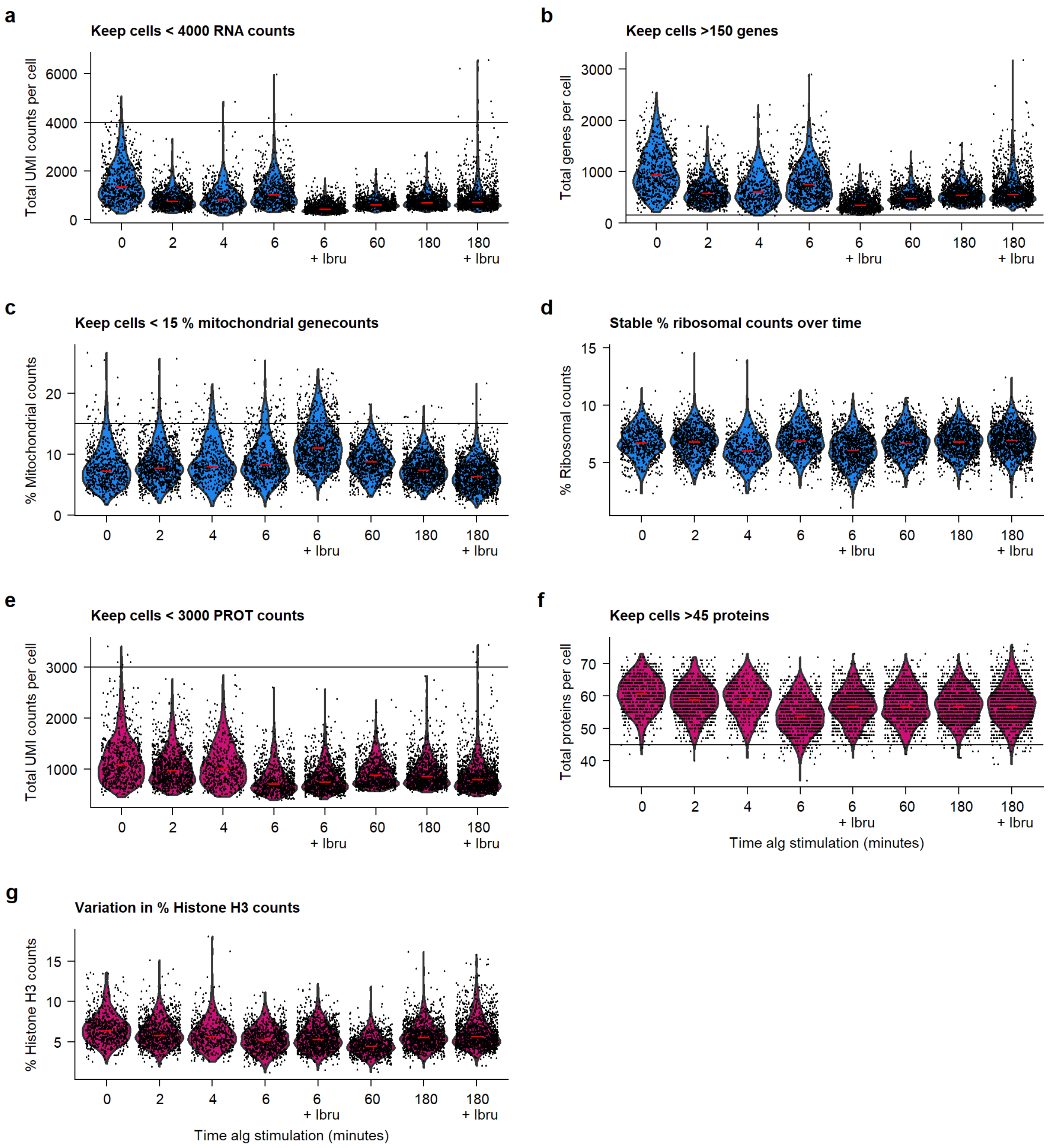


**Supplementary Figure 6: Technical quality of the QuRIE-seq transcriptomic and proteomic libraries and quality checks filtering for fixed BJAB cells at eleven different stimulation and/or inhibition conditions: after stimulation with alg for: 0, 2, 4, 6, 60 and 180 minutes, with or without prior Ibrutinib inhibition resulted in a multimodal dataset of 6952 cells with matched gene and (phospho)protein expression levels.** Violin plots for (**a-d, blue**) the transcriptomic and (**e-g, pink**) the proteomic libraries. **a**, The number of unique molecular identifiers filtered mapped (UMIFM) detected per cell. Cells with < 4000 UMIFM counts per cell were kept for further analysis. **b**, The number of different genes detected per cell. Cells with > 150 genes per cell were kept for further analysis. **c**, Percentage of mitochondrial UMIFM counts out of the total number of UMIFM counts per cell. Cells with < 15% mitochondrial counts per cell were kept for further analysis. **d**, Percentage of ribosomal UMIFM counts out of the total number of UMIFM counts per cell. **e**, The number of protein counts per cell. Cells with < 3000 protein counts per cell were kept for further analysis. **f**, The number of different proteins detected per cell. Cells with > 45 different proteins detected were kept for further analysis. **g**, The percentage of Histone H3 counts out of the total number of protein counts per cell.

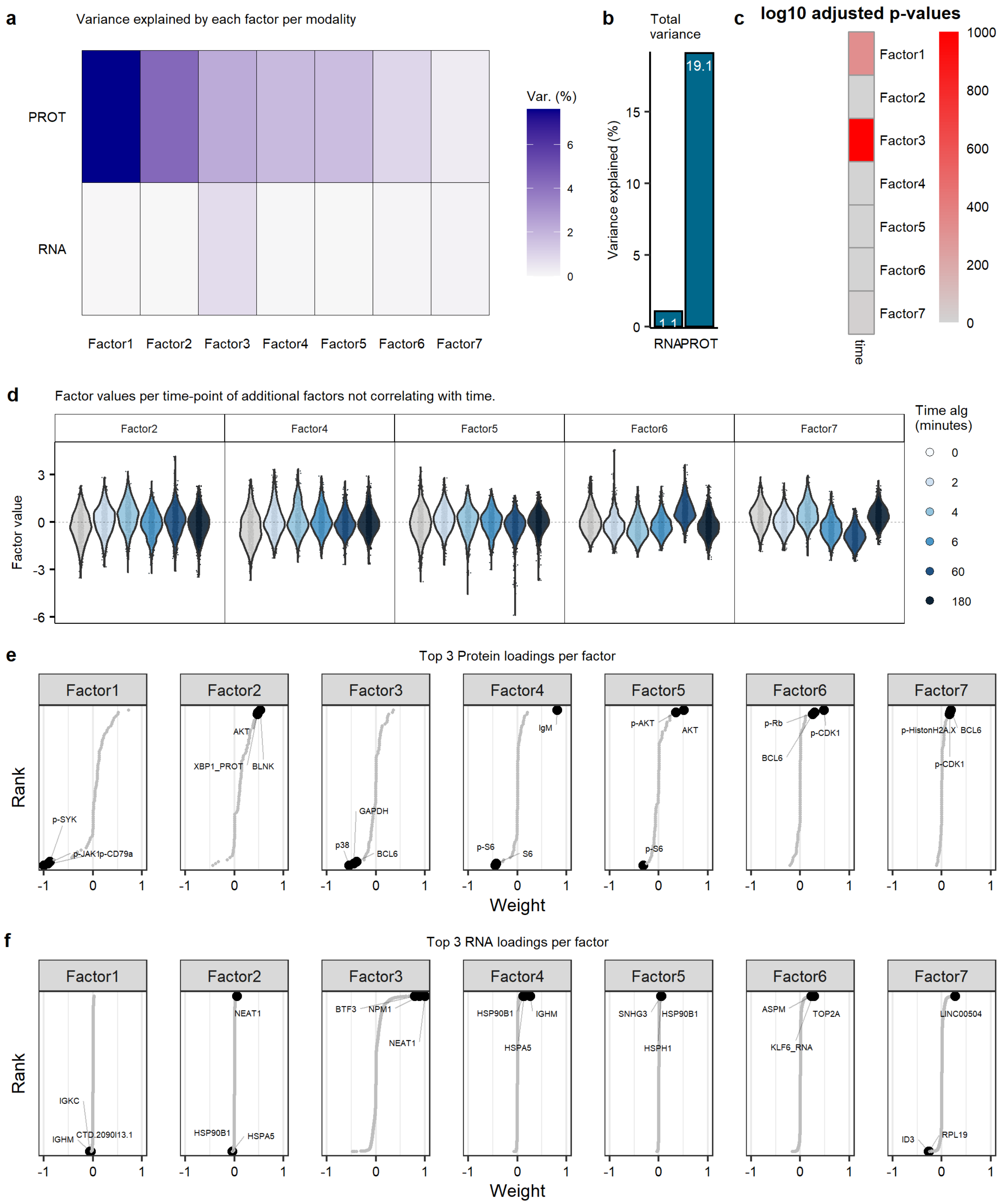


**Supplementary Figure 7: Supplementary results on MOFA+ analysis Fig. 2. a**, Percentage of variance explained by each factor per modality (proteins and RNA). **b**, Percentage of the total variance explained per modality. **c**, Pearson's correlation (p-value) between the MOFA+ factors and the duration of anti-immunoglobulin antibody (aIg) stimulation. **d**, Violin plots of the factor values as a function of aIg stimulation for: 0, 2, 4, 6, 60 and 180 minutes. Factor 1 and 3 are shown in Fig. 2a,e respectively. **e**, Protein loadings per factor. The names of the top three proteins loading per factor are marked in the plots. **f**, RNA loadings per factor. The name of the top three RNA loadings per factor are marked in the plots.

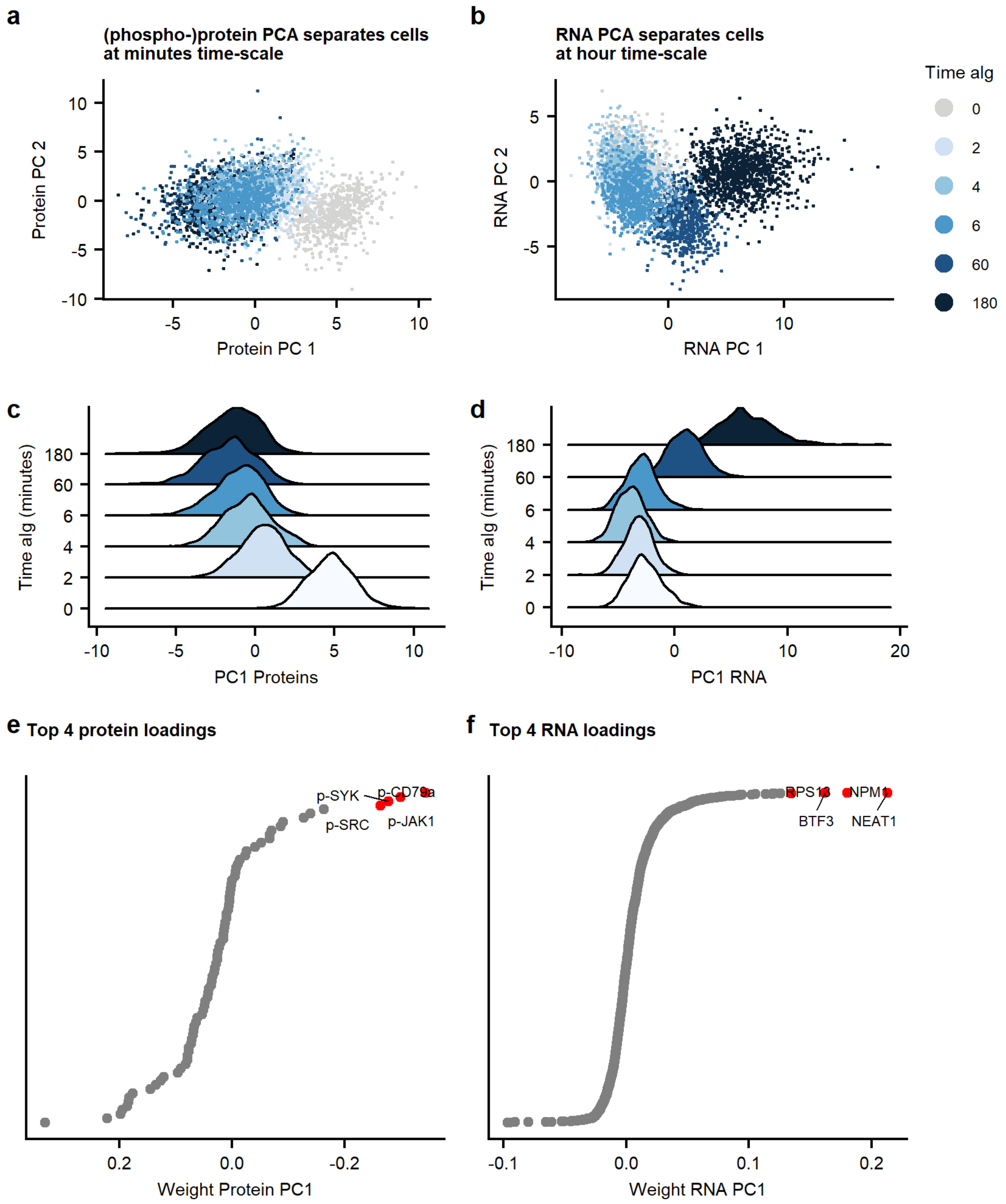


**Supplementary Figure 8: PCA on protein and RNA dataset shows the equivalence of PC1 in separate modalities: proteomics and transcriptomics to factor 1 and factor 3 respectively in MOFA+ analysis.** **a**, Principal component analysis (PCA) plot (PC1 and PC2) using the proteomic data. **b**, PCA plot (PC1 and PC2) using the transcriptomic data. **c**, PC1 as a function of the duration of αIg stimulation for the proteomic data. **d,** PC1 as a function of the duration of αIg stimulation for the transcriptomic data. **e,** Protein loadings contributing to PC1. The top 4 protein loadings are annotated. **f**, RNA loadings contributing to PC1. The top 4 RNA loadings are annotated. The results are equivalent to factors 1 and 3 in MOFA+ analysis, respectively.

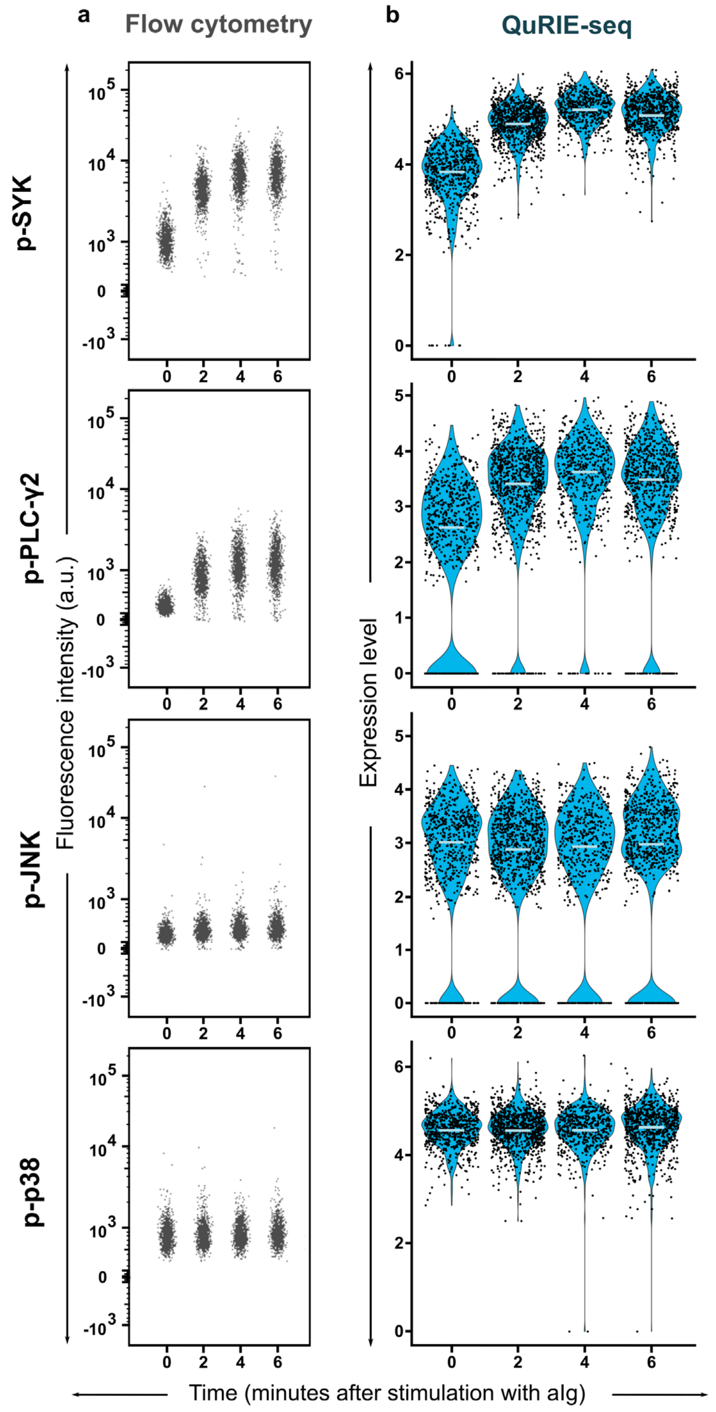


**Supplementary Figure 9**: **Regulation of p-SYK, p-PLC-γ2, p-JNK and p-p38 protein expression upon stimulation with aIg in BJAB cells.** The phosphorylation level of four proteins was determined using: **a**, flow cytometry, and **b**, QuRIE-seq technology. Different durations of stimulation were compared: 0, 2, 4 and 6 minutes.

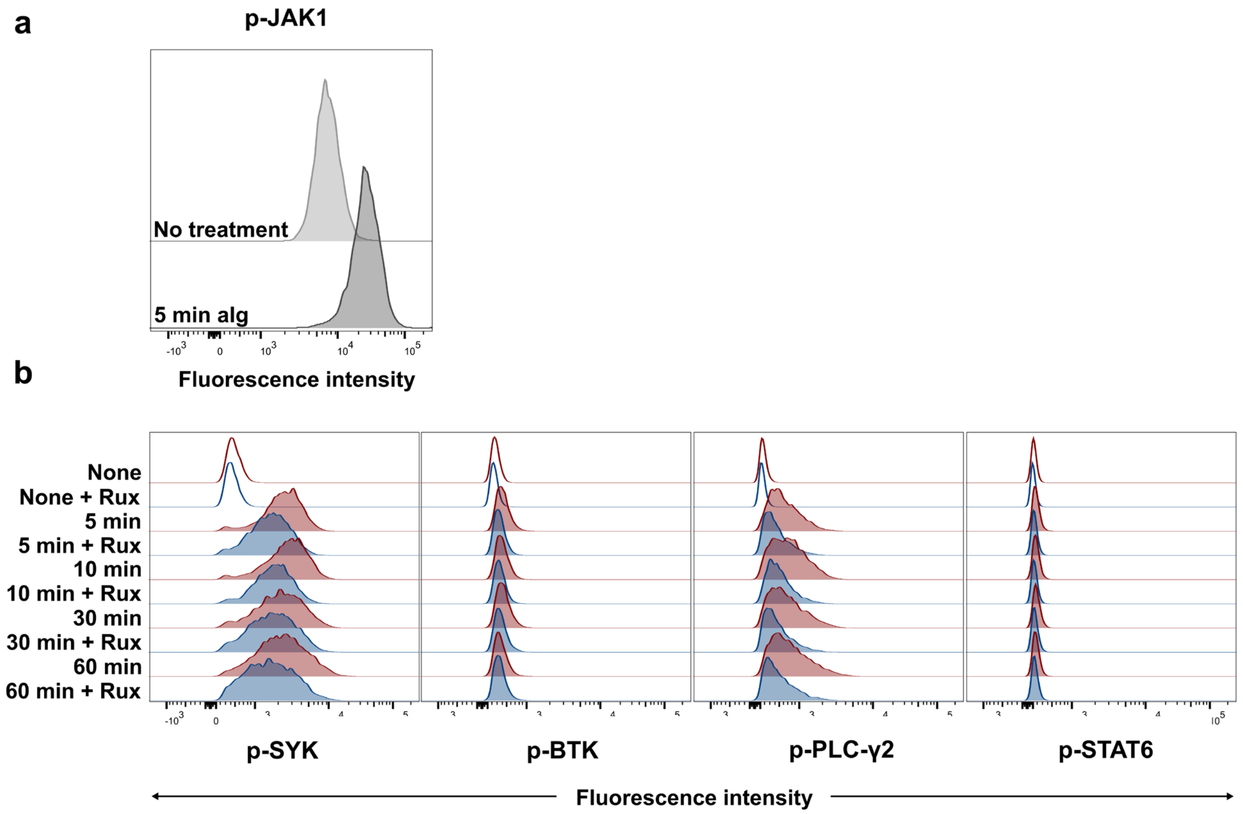


**Supplementary Figure 10: The role of JAK1 in the BCR response to stimulation with aIg. a,** JAK1 phosphorylation after 5 minutes stimulation with aIg characterized by flow cytometry. **b**, The phosphorylation level of four proteins: SYK, BTK, PLC-γ2, and STAT6 determined using flow cytometry for unstimulated BJAB cells, and stimulated with aIg for: 5, 10, 30, and 60 minutes with (blue curves) or without (red curves) previous Ruxolitinib (Rux) inhibition. *Rux is a JAK1/2 inhibitor.*

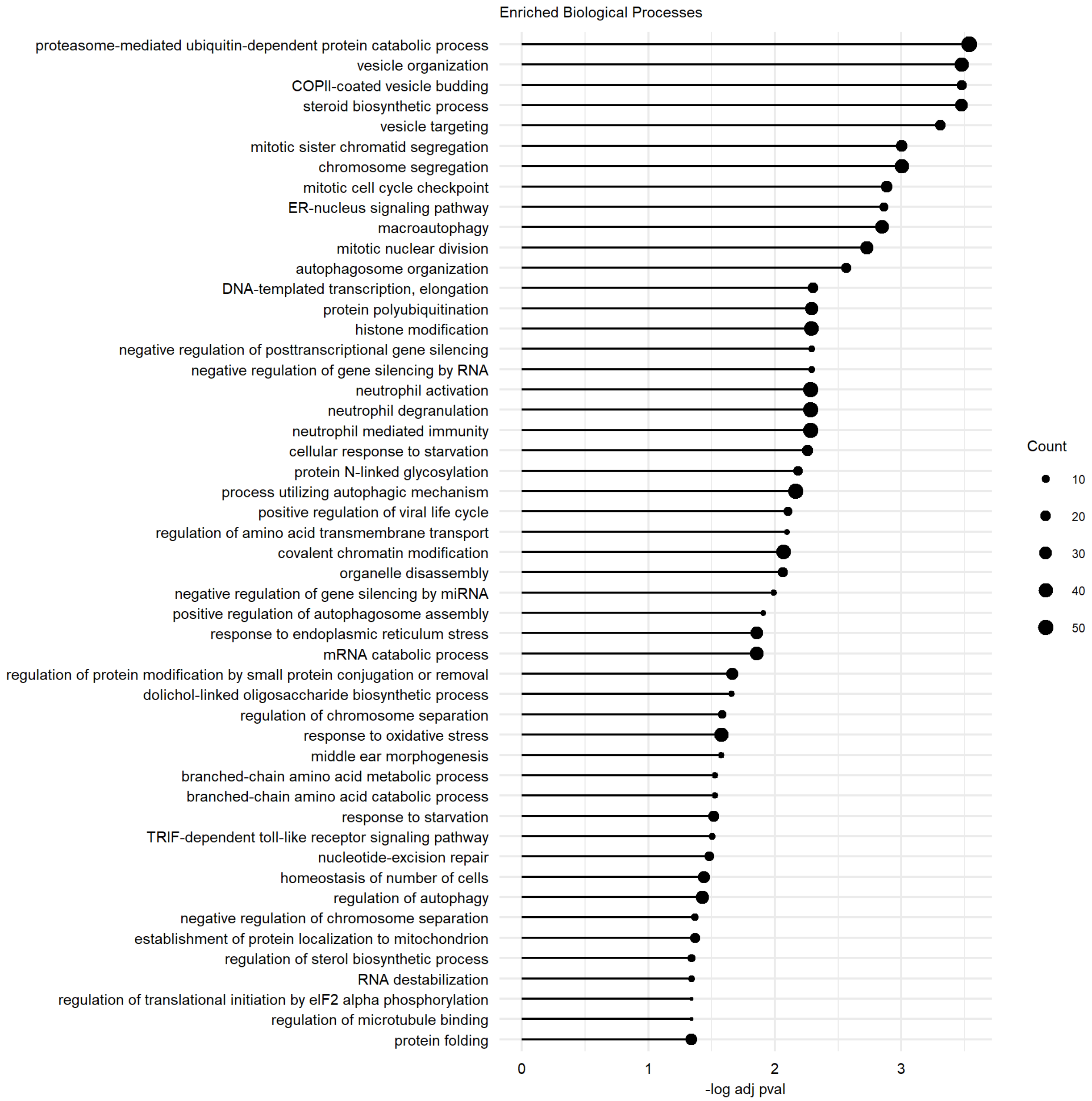


**Supplementary Figure 11: Top 50 gene-set (biological processes) significantly enriched in genes with positive factor 3 loadings.**

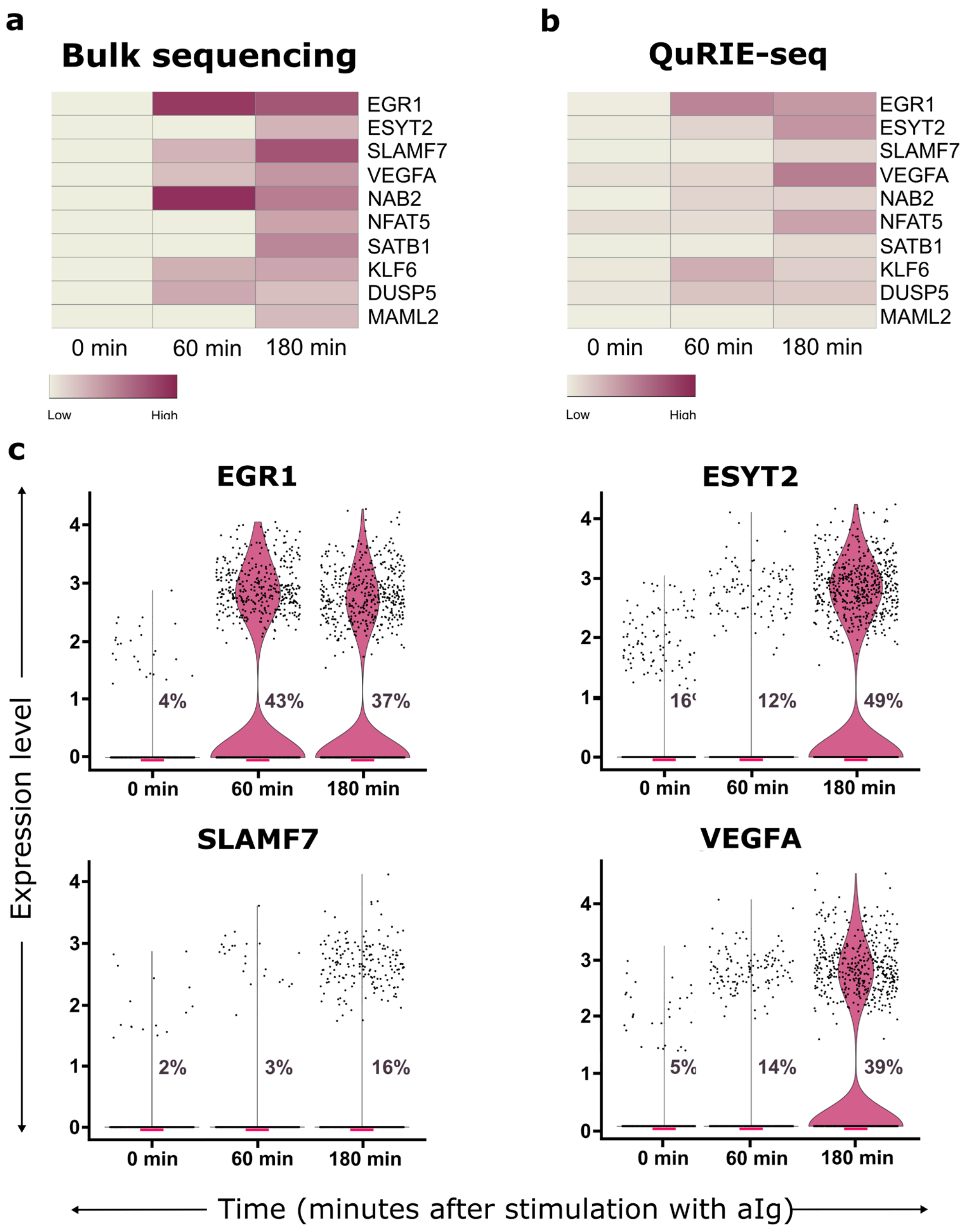


**Supplementary Figure 12**: **Regulation of gene expression in BJAB cells upon stimulation with aIg.** The expression of ten genes (*EGR1*, *ESYT2*, *SLAMF7*, *VEGFA*, *NAB2*, *NFAT5*, *SATB1*, *KLF6*, *DUSP5*, *MAML2*) was determined using: **a**, bulk sequencing, and **b**, the individual cell expression (determined by QuRIE-seq) averaged over the whole population, and plotted as a heatmap. **c**, The expression of each single-cell determined by QuRIE-seq measurement is shown as a violin plot for four different genes: *EGR1*, *ESYT2*, *SLAMF7*, and *VEGFA*; the percentages indicate the fraction of cells with the gene expression >0 for each stimulation time point (0, 60, 180 minutes).

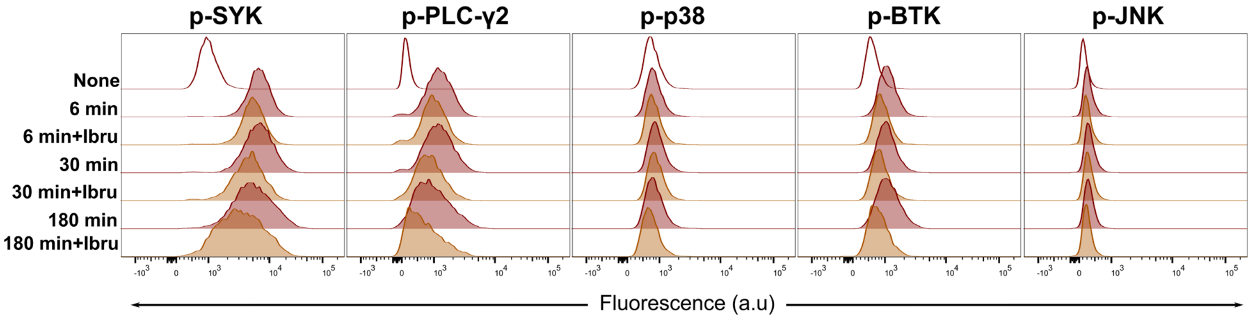


**Supplementary Figure 13: Validation by flow cytometry of inhibition with Ibrutinib (Ibru) of the BCR pathway.** The phosphorylation of SYK, PLC-γ2, p38, BTK, and JNK was characterized upon different stimulation conditions: unstimulated cells (0 minutes - red empty line); stimulation with aIg for: 6, 30, and 180 minutes with (orange filled line) and without (red filled line) prior Ibru inhibition.

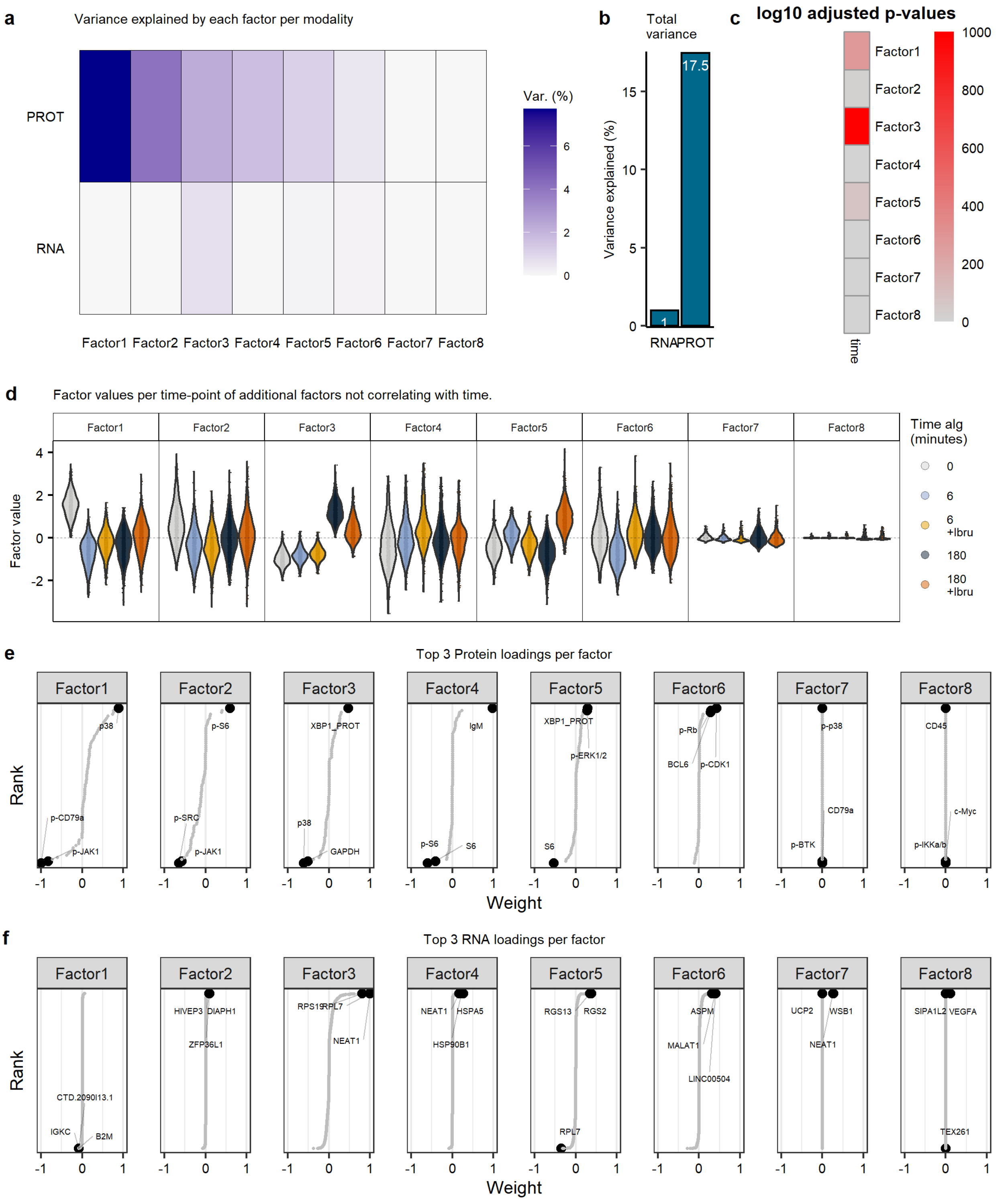


**Supplementary Figure 14: Supplementary results on MOFA+ analysis Fig. 3. a**, Percentage of variance explained by each factor per modality (proteins and RNA). **b**, Percentage of the total variance explained per modality. **c**, Pearson's correlation (p-value) between the MOFA+ factors and the duration of anti-immunoglobulin antibody (aIg) stimulation. **d**, Violin plots of the factor values as a function of stimulation with aIg for: 0, 6, and 180 minutes with and without Ibru. **e**, Protein loadings per factor. The names of the top three proteins loading per factor are marked in the plots. **f**, RNA loadings per factor. The name of the top three RNA loadings per factor are marked in the plots.

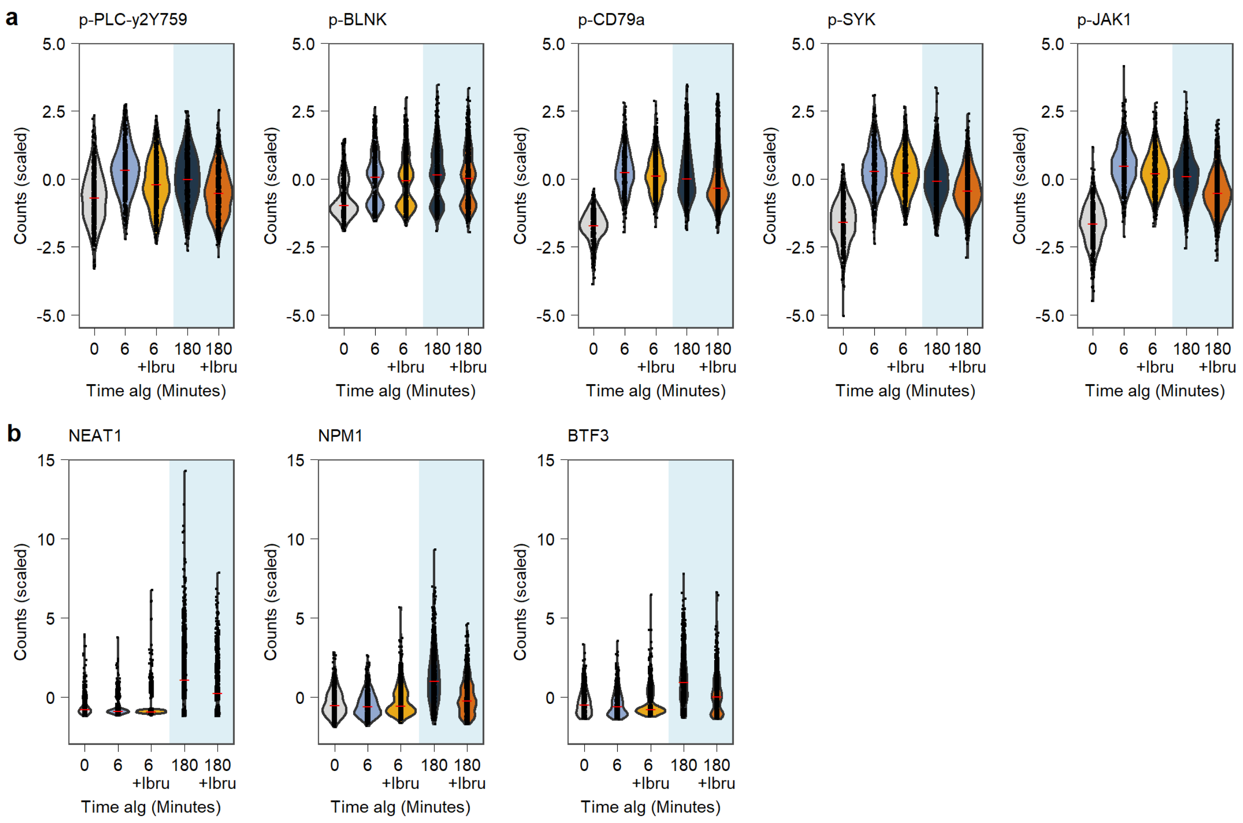


**Supplementary Figure 15: Phosphorylation up- and downstream of BTK, and downstream genes, dampened by Ibrutinib (Ibru) measured with QuRIE-seq. a**, Phosphoprotein scaled counts of signaling components up- or downstream of BTK: PLC-y2, BLNK, CD79a, SYK, and JAK1 as a function of stimulation with aIg for: 0, 6, and 180 minutes with and without Ibru. **b**, Top 3 genes of factor 3 loadings: *NEAT1*, *NPM1*, and *BTF3*, scaled counts as a function of stimulation with aIg for: 0, 6, and 180 minutes with and without Ibru.

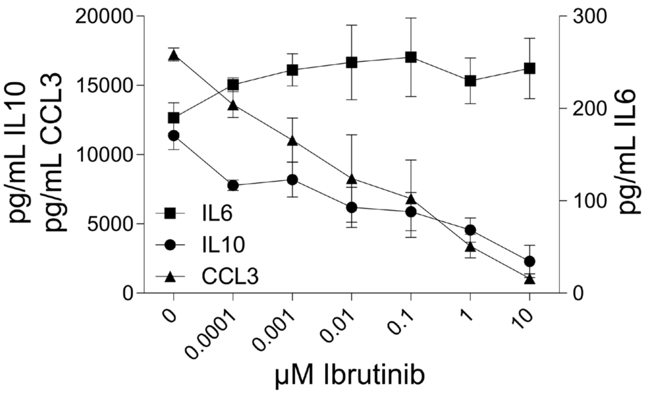


**Supplementary Figure 16: Analysis of BJABs culture supernatant for cytokines upon overnight (16 hours) stimulation in presence of Ibrutinib.** The BTK inhibitor (Ibrutinib) was used to block BTK (part of BCR signaling pathway), and titrated while cells were stimulated with 10 µg/ml aIg. The dose-dependent decrease of cytokine secretion was observed through ELISA for IL10 and CCL3, but not for IL6.

#### Supplementary Tables

**Supplementary Table 1: Antibodies used for immunostaining of cells prior to microfluidic encapsulation.**

| | Target | Ab barcode sequence | Vendor | Clone | Used concentration |
|---|---|---|---|---|---|
| 1 | **Histone H3** | **CAATCCCT** | Cell Signaling 4499BF | D1H2 | 0.1 µg/ml |
| 2 | **p-p65** | **GTCCAGGC** | Cell Signaling 3033BF | 93H1 | 1 µg/ml |
| 3 | **p-SYK** | **TGTGTATA** | Cell Signaling 2710BF | C87C1 | 1 µg/ml |
| 4 | **p-JNK** | **AGGATCGA** | Cell Signaling 9255BF | G9 | 1 µg/ml |
| 5 | **p-p38** | **CACGATTC** | Cell Signaling 4511BF | D3F9 | 1 µg/ml |
| 6 | **p-PLC-y2** | **GTATCGAG** | R&D Systems MAB37161 | 790623 | 1 µg/ml |
| 7 | **p-BTK** | **TCTCGACT** | Biolegend 601702 | A16128B | 1 µg/ml |
| 8 | **p-SHP-2** | **ACCCGCAC** | Cell Signaling 5431BFF | D66F10 | 1 µg/ml |
| 9 | **p-SHP-1** | **CATGCGTA** | Cell Signaling 8849BF | D11G5 | 1 µg/ml |
| 10 | **Cyclin E** | **GTGATAGT** | ThermoFisher 32-1600 | HE12 | 1 µg/ml |
| 11 | **c-MYC** | **TGATATCG** | ThermoFisher 700648 | 27H46L35 | 1 µg/ml |
| 12 | **p-RB** | **AGGGCGTT** | ThermoFisher 701059 | 14H7L14 | 0.1 µg/ml |
| 13 | **p-c-JUN** | **CTATACGC** | ThermoFisher MA5-27760 | GT653) | 1 µg/ml |
| 14 | **active Caspase-3** | **GCTCGTCA** | BD Bioscience 559565 | C92-605 | 1 µg/ml |
| 15 | **CD70** | **TACATAAG** | BD Bioscience 555833 | Ki-24 | 1 µg/ml |
| 16 | **IgD** | **AATTGAAC** | BD Bioscience 555776 | IA6-2 | 1 µg/ml |
| 17 | **CD86** | **CCAGTGGA** | BD Bioscience 555655 | 2331 (FUN-1) | 1 µg/ml |
| 18 | **CD20** | **GTCCATTG** | BD Bioscience 555677 | H1 | 0.1 µg/ml |
| 19 | **CD79a** | **TGGACCCT** | BD Bioscience 555934 | HM47 | 0.1 µg/ml |
| 20 | **CD5** | **AGCAGTTA** | BD Bioscience 555350 | UCHT2 | 1 µg/ml |
| 21 | **p-AKT** | **CTTGTACC** | Cell Signaling 4060BF | D9E | 1 µg/ml |
| 22 | **S6** | **GAACCCGG** | Cell Signaling 2317B | 54D2 | 1 µg/ml |
| 23 | **ERK 1/2** | **TCGTAGAT** | Cell Signaling 4696BF | L34F12 | 1 µg/ml |
| 24 | **p38** | **ACGCGGAA** | Cell Signaling 8690BF | D13E1 | 1 µg/ml |
| 25 | **AKT** | **CGCTATCC** | Cell Signaling 4685BF | 11EE7 | 1 µg/ml |
| 26 | **SYK** | **GTTGCATG** | Cell Signaling 13198BF | D3Z1E | 1 µg/ml |
| 27 | **BTK** | **TAAATCGT** | Cell Signaling 8547BF | D3H5 | 0.1 µg/ml |
| 28 | **p-S6** | **ATCGCCAT** | Cell Signaling 4858BF | D57.2.2E | 0.1 µg/ml |
| 29 | **p-ERK 1/2** | **CATAAAGG** | Cell Signaling 5726BF | D1H6G | 1 µg/ml |
| 30 | **p65** | **TCACGGTA** | Cell Signaling 8242BF | D14E12 | 0.1 µg/ml |
| 31 | **CD19** | **CACTCAAC** | SantaCruz sc-373897 | F-3 | 2 µg/ml |
| 32 | **p-Histon H3** | **GCTGTGA** | Biolegend 650802 | 11D8 | 0.1 µg/ml |
| 33 | **IgM** | **TTGCGTCG** | Biolegend 314502 | MHM-88 | 1 µg/ml |
| 34 | **Cyclin A** | **ATATGAGA** | Biolegend 644001 | E23.1 | 0.1 µg/ml |
| 35 | **p-Histon H2A.X** | **CACCTCAG** | Biolegend 613402 | 2F3 | 0.1 µg/ml |
| 36 | **Cyclin B1** | **GCTACTTC** | Biolegend 647902 | V152 | 0.1 µg/ml |

| | | | | | |
|---|---|---|---|---|---|
| **37** | **Ki-67** | **TGGGAGCT** | Biolegend 350523 | Ki-67 | 0.1 µg/ml |
| **38** | **JNK** | **ATCCGGCA** | R&D Systems AF1387 | | 1 µg/ml |
| **39** | **SHP-1** | **CCGTTATG** | R&D Systems MAB1878 | 255402 | 1 µg/ml |
| **40** | **p-SRC** | **GGTAATGT** | R&D Systems MAB2685 | 1246F | 1 µg/ml |
| **41** | **p-TOR** | **TAAGCCAC** | R&D Systems MAB1665 | 834115 | 0.1 µg/ml |
| **42** | **GAPDH** | **ACCGAACA** | Biolegend 607902 | W17079A | 1 µg/ml |
| **43** | **BLNK** | **GTTTGTGG** | BD Biosciences 559930 | 2B11 | 1 µg/ml |
| **44** | **p-BLNK** | **TAGACGAC** | BD Biosciences 558366 | J117-1278 | 0.1 µg/ml |
| **45** | **p-PKC-b1** | **ACGCTTGG** | ThermoFisher 702430 | 3H8L1 | 0.1 µg/ml |
| **46** | **CD53** | **CGCTACAT** | BD Biosciences 555506 | HI29 | 0.1 µg/ml |
| **47** | **CD38** | **GAAAGACA** | Biolegend 303535 | HIT2 | 0.1 µg/ml |
| **48** | **CD45** | **TTTGCGTC** | Biolegend 304045 | HI30 | 0.1 µg/ml |
| **49** | **p-IRAK4** | **ATGGTCGC** | Cell Signaling 11927S | D6D7 | 1 µg/ml |
| **50** | **p-CD79a** | **CGACATAG** | Cell Signaling 14732BF | D1B9 | 0.1 µg/ml |
| **51** | **p-CDK1** | **GATTCGCT** | ThermoFisher 701808 | 17H29L7 | 1 µg/ml |
| **52** | **p-CDK4** | **TCCAGATA** | ThermoFisher 702556 | 9H2L7 | 1 µg/ml |
| **53** | **p-AMPK-a1/2** | **ACTACTGT** | ThermoFisher701068 | 10H2L20 | 0.1 µg/ml |
| **54** | **p-AMPK-b1** | **CGGGAACG** | Thermo Fisher 700241 | 9H26L42 | 0.1 µg/ml |
| **55** | **T-bet** | **GACCTCTC** | Biolegend 644825 | 4B10 | 1 µg/ml |
| **56** | **p-IKK a/b** | **TTATGGAA** | ThermoFisher 701643 | 7H17L17 | 1 µg/ml |
| **57** | **CD27** | **ACAGCAAC** | Abcam ab192336 | EPR8569 | 0.1 µg/ml |
| **58** | **p-JAK1** | **CGCAATTT** | ThermoFisher 700028 | 59H4L5 | 1 µg/ml |
| **59** | **p-PLC-γ2 (Y759)** | **GAGTTGCG** | R&D Systems MAB7377 | 744757 | 0.1 µg/ml |
| **60** | **IL10** | **TTTCGCGA** | ThermoFisher 16-7108-85 | JES3-9D7 | 1 µg/ml |
| **61** | **CCL3/4** | **ACCAGTCC** | R&D Systems MAB2701-100 | 93342 | 1 µg/ml |
| **62** | **CD80** | **CTTTCCTT** | Biolegend 305212 | 2D10 | 1 µg/ml |
| **63** | **p-CDK6** | **CTGGACGT** | ThermoFisher | 16HCLC | 1 µg/ml |
| **64** | **p-STAT1** | **GAACGGTC** | ThermoFisher 33-3400 | ST1P-11A5 | 1 µg/ml |
| **65** | **p-STAT3** | **TGTTCACG** | Biolegend 690402 | A16002B | 1 µg/ml |
| **66** | **p-STAT5** | **ATCTGATC** | ThermoFisher 701063 | 6H5L15 | 1 µg/ml |
| **67** | **p-STAT6** | **GAGAAGGG** | ThermoFisher 700247 | 46H1L12 | 1 µg/ml |
| **68** | **KLF6** | **TCACTCCT** | ThermoFisher 39-6900 | 9A2 | 1 µg/ml |
| **69** | **BCL6** | **AGATAACA** | BD Bioscneces 561520 | K112-91 | 1 µg/ml |
| **70** | **AID** | **CTTATTTG** | BD Biosciences 565784 | EK2-5G9 | 1 µg/ml |
| **71** | **IgG** | **GCGGGCAT** | BD Biosciences 555784 | G18-145 | 0.1 µg/ml |
| **72** | **CD24** | **TACCCGGC** | RnD Systems MAB5247 | ML5 | 1 µg/ml |
| **73** | **IgA** | **ATTGTTTC** | Biolegend 411502 | HP6123 | 0.5 µg/ml |
| **74** | **IgE** | **CGCAGGAG** | Biolegend 325502 | MHE-18 | 1 µg/ml |
| **75** | **CD23** | **GCACCAGT** | Abcam ab245732 | SP163 | 0.1 µg/ml |
| **76** | **BLIMP1** | **TAGTACCA** | RnD Systems MAB36081 | 646702 | 1 µg/ml |

| | | | | | |
|---|---|---|---|---|---|
| **77** | **IRF8** | **ATTCGTGC** | Biolegend 656502 | 656502 | 1 µg/ml |
| **78** | **IRF4** | **CGCGTGCA** | ThermoFisher 14-9858-82 | 3 E 4 | 1 µg/ml |
| **79** | **BAFF-R** | **GAATACTG** | RnD Systems MAB1162 | 2403C | 1 µg/ml |
| **80** | **XBP1** | **TCGACAAT** | Abcam ab239954 | EPR4086 | 1 µg/ml |

**Supplementary Table 2: Antibodies used for flow cytometry analysis of BJABs.**

| **Target** | **Fluorophore** | **Vendor** | **Clone** | **Used concentration** |
|---|---|---|---|---|
| **p-SYK** | PE | Cell Signaling 6485 | C87C1 | 1/50 |
| **p-PLC-y2** | Alexa 647 | BD Bioscience 558498 | K86-689.37 | 1/100 |
| **p-BTK** | PE | Biolegend 601704 | A16128B | 1/50 |
| **p-p38** | Alexa 488 | Cell Signaling 41768S | 3D7 | 1/50 |
| **p-AKT** | PE | Cell Signaling 5315 | D9E | 1/50 |
| **p-ERK 1/2** | PE | Cell Signaling 75765S | D1H6G | 1/50 |
| **p-JNK** | Alexa 647 | Cell Signaling 9257S | G9 | 1/50 |
| **p-p65** | Alexa 647 | Cell Signaling 5733 | 93H1 | 1/50 |
| **p-S6** | PE | Cell Signaling 5316 | D57.2.2E | 1/800 |
| **CD19** | PerCP/Cy5.5 | Biolegend 302230 | HIB19 | 1/100 |
| **IgD** | BV510 | Biolegend 348220 | IA6-2 | 1/50 |
| **CD27** | FITC | BD Bioscience 555440 | M-T271 | 1/25 |
| **aCD20** | APC-H7 | BD Bioscience 560853 | 2H7 | 1/25 |
| **CD38** | PE-Cy7 | eBioscience 250-388-41 | HB7 | 1/2000 |
| **IgA** | PE | Miltenyi 130-113-472 | IS11-8E10 | 1/200 |

**Supplementary Table 3: Sequences of all the primers used for library preparation.** All primers were ordered from Biolegio (The Netherlands).

| | Primer sequence 5’ -> 3’ | Length (bp) |
|---|---|---|
| **PE2-N6** | TCGGCATTCCTGCTGAACCGCTCTTCCGATCT NNNNN | 38 |
| **PE1** | CAAGCAGAAGACGGCATACGAGAT [6-bp library index] CTCTTTCCCTACACGA | 46 |
| **PE2** | AATGATACGGCGACCACCGAGATCTACACGGTCTCGGCATTCCTGCTG AAC | 51 |
| **PE2-NNNN-Next1** | TCGGCATTCCTGCTGAACCGCTCTTCCGATCTNNNNGATGTGTATAAG AGAC | 52 |
| **PE2-NNNN-BioHash2** | TCGGCATTCCTGCTGAACCGCTCTTCCGATCTNNNNCGTGTGCTCTTCC GAT | 52 |
| **PE2-Nextera2 (long)** | AATGATACGGCGACCACCGAGATCTACACGGTCTCGTCGGCAGCGTCA GATG | 52 |
| **Custom Read 1 primer** | GGCATTCCTGCTGAACCGCTCTTCCGATCT | 30 |
| **Custom Index Read primer** | AGATCGGAAGAGCGTCGTGTAGGGAAAGAG | 30 |
| **Custom Read 2 primer** | CTCTTTCCCTACACGACGCTCTTCCGATCT | 30 |
